# Nondestructive Spatial Lipidomics for Glioma Classification

**DOI:** 10.1101/2023.03.09.531882

**Authors:** Davide Alessandro Martella, Leor Ariel Rose, Nadia Rouatbi, Chenlei Gu, Valeria Caprettini, Magnus Jensen, Cathleen Hagemann, Andrea Serio, Khuloud Al-Jamal, Maddy Parsons, Mads Bergholt, Paul Brennan, Assaf Zaritsky, Ciro Chiappini

## Abstract

Mapping the molecular composition of tissues using spatial biology provides high-content information for molecular diagnostics. However, spatial biology approaches require invasive procedures to collect samples and destroy the investigated tissue, limiting the extent of analysis, particularly for highly functional tissues such as those of the brain. To address these limitations, we developed a workflow to harvest biomolecules from brain tissues using nanoneedles and characterise the distribution of lipids using desorption electrospray ionization mass spectrometry imaging. The nanoneedles preserved the original tissue while harvesting a reliable molecular profile and retaining the original lipid distribution for mouse and human brain samples, accurately outlining the morphology of key regions within the brain and tumour lesions. The deep neural network analysis of a cohort containing 23 human glioma biopsies showed that nanoneedle samples maintain the molecular signatures required to accurately classify disease state. Thus, nanoneedles provide a route for tissue-preserving spatial lipidomic and molecular diagnostics.

---

Spatial biology provides-omics level information on the molecular landscape of complex systems such as tissues and organotypic models and contributes to investigating regulatory networks within microenvironments and the non-cell autonomous processes underpinning health, disease, and development. In particular, mapping the molecular composition of tumours can elucidate the interactions underlying their progression and heterogeneity, improving classification for treatment selection. Indeed, spatial transcriptomics, proteomics, and metabolomics have helped elucidate the tumour microenvironment and its role in disease progression and invasion ^1–3^.

Spatial biology approaches require tissue sections acquired through invasive procedures such as biopsies and employ processes that largely sacrifice the specimen. In cancer diagnostics, the need to minimise invasiveness and to preserve organ function limits the number, frequency and extent of samples available, posing a challenge in acquiring an accurate spatial molecular characterisation of the lesions, particularly hampering longitudinal analysis and the investigation of non-excisable tumours.

Vertical nanoprobes such as nanoneedles could provide a route for minimally invasive collection, to enable sampling more tissue and performing multiple analyses on the same specimen. Vertical nanoprobes are high-aspect nanomaterials capable of probing and manipulating the intracellular space with minimal perturbation^4,5^. Nanoprobes interfaced with living tissues can harvest biomolecules without impacting cell function^6^. Since the process is nonperturbing, it can be repeated frequently without significant invasiveness. Leveraging this non-perturbing approach, atomic force microscopy (AFM)-operated nanoprobes can repeatedly sample the cytosolic fluid of the same cells, and monitor the evolution of the transcriptional landscape in response to stimuli^7^. Arrays of nanoprobes decorated with ligands can repeatedly capture individual analytes or small classes of biomolecules for direct molecular recognition to register temporal variations in signalling processes of living cell cultures and tissues^8^. In cancer diagnostics, nanoprobes can characterise activity levels of individual biomarkers across human tissue samples with cellular resolution, mapping tissue heterogeneity^9^.

These results highlight the yet unfulfilled potential of vertical nanoprobes to harvest and analyse a *molecular replica*, defined as a spatially-preserved, representative sample of the biomolecules within a tissue, while preserving the original sample.

Developing an approach for minimally invasive molecular diagnostics using a molecular replica could benefit the treatment of gliomas. Glioma is a primary malignant brain tumour which can reprogram lipid metabolism to promote cell membrane integrity and support abnormal cell proliferation^10^. Aberrant lipid profiles are exploited by malignant cells to thrive and are appealing targets for novel therapies. Heterogeneous lipid dysregulation can carry relevant information for clinical intervention^11,12^, highlighting the value to explore their spatial distribution. However, acquiring the necessary brain tissue samples is hampered by the high risk of injury associated with biopsy sample collection. As limited tissue samples are available, molecular replicas could improve lipidomic mapping of gliomas, refining diagnosis and therapy.

Desorption electrospray ionisation mass spectrometry imaging (DESI-MSI) is a widely adopted approach to map lipids and metabolites. DESI-MSI of human brain samples can investigate remyelination processes in multiple sclerosis^13^. Applied to glioma diagnostics, it can grade tumours with 97.9 % accuracy, and classify IDH mutant status based on metabolic and lipid profile^14^.

Here we developed an integrated workflow for comprehensive characterisation of human gliomas by spatial lipidomics on molecular tissue replicas generated with nanoneedle arrays (Figure 1). In this workflow we applied a nanoneedle array to brain tissue, harvesting a molecular replica. Analysis of the molecular replica by DESI-MSI provided a lipidomic map capable of discriminating key brain morphological features, in agreement with those of the original tissue with respect to the relative abundance of species and their spatial distribution. The analysis of these maps through deep neural networks for a set of high and low grade human gliomas showed a comparable classification power for molecular replicas and tissue sections.

**Figure 1:**
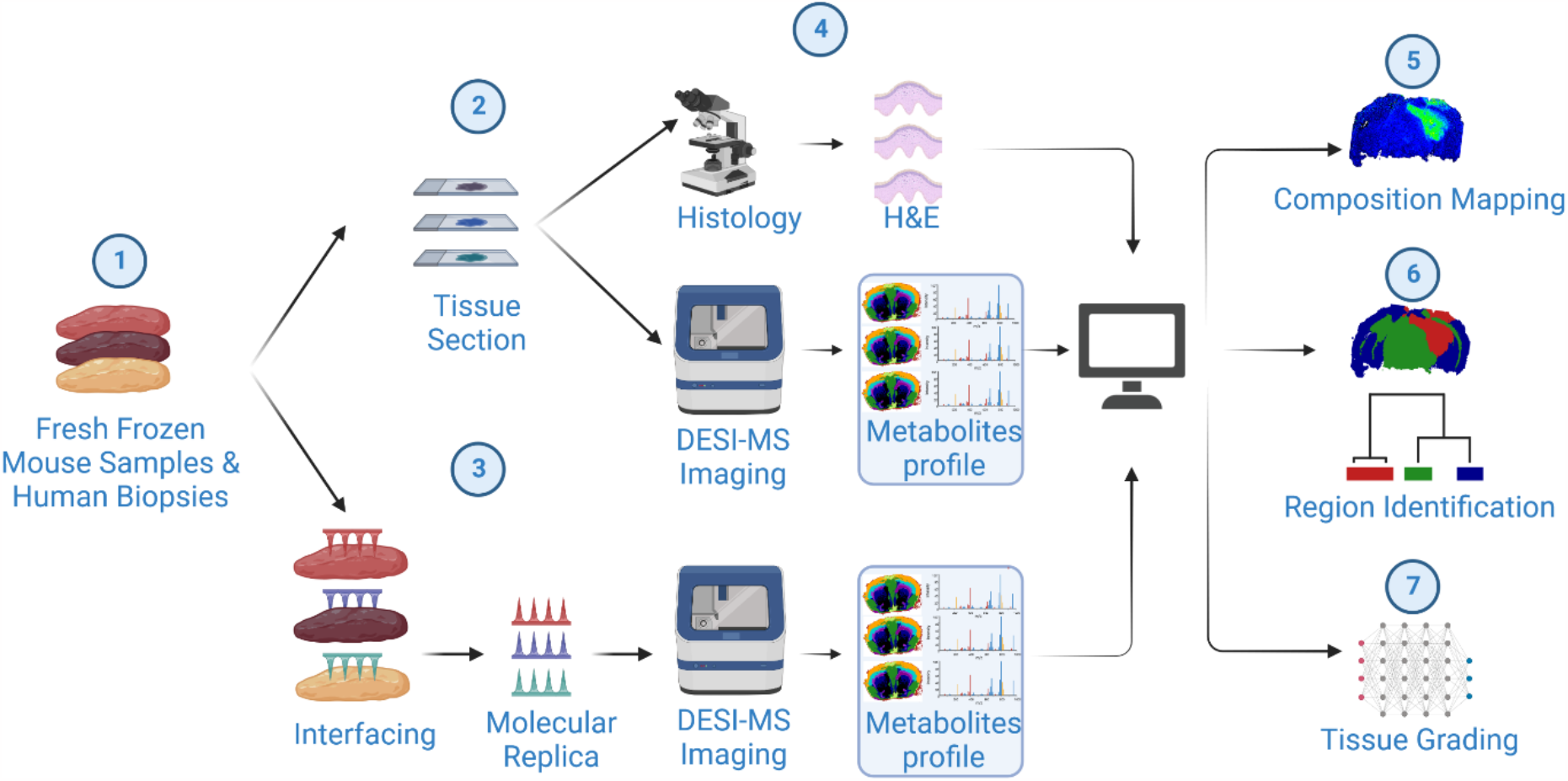
The molecular replica workflow. Mouse brain and human glioma biopsies (1) are sectioned to obtain reference tissue samples (2), and interfaced with nanoneedles to generate molecular replicas (3). For each sample we generated at least one DESI-MSI dataset from a tissue section proximal to the molecular replica to obtain a metabolite reference map and at least one H&E dataset to obtain a morphological reference (4). We analysed the DESI-MSI datasets of section and replica to assess their relative performance. We compared the intensity distribution of key lipids to map the tissue composition(5); the ability to outline tissue regions such as white matter, grey matter and different types of lesions (6), and the ability to determine tissue similarity and grade from human glioma biopsies (7).

## Results

### Using nanoneedles to generate molecular replicas

We first evaluated the ability of nanoneedles formed on a silicon chip to generate a molecular replica from fresh frozen mouse brains by comparing them to an unprocessed, flat silicon chip (flat substrate). In order to generate the replica we interfaced an 8×8 mm^2^ square chip with the frozen tissue. Computational modelling of the interfacing indicated that, upon contact, a few microns of tissue transiently thaws, allowing harvesting of biomolecules (Figure S1). We designed an interfacing device to minimise lateral movement and self-align the substrate with the tissue, ensuring stable and conformal contact (Figure 2a,b). Following interfacing, scanning electron microscopy showed a uniform molecular adsorbate on the nanoneedles, while the flat substrate lacked uniformity showing areas without coverage (holes) and areas of adsorbate accumulation (lumps) (Figure 2c). Fluorescence microscopy following lipid staining showed a nanoneedle replica fully reproducing the original tissue and macroscopically retaining its morphology (Figure 2d). The flat replica only reproduced a small portion of the original tissue which lacked a clear morphology within the harvested lipids. While the nanoneedle replicated the entire tissue area, the area of the flat replica amounted to less than half of that of the reference tissue (Figure 2e). These data indicated that, unlike flat substrates, nanoneedles can generate uniform molecular replicas for lipidomic analysis by sampling the entirety of the interfacing tissue and retaining its original morphology.

**Figure 2:**
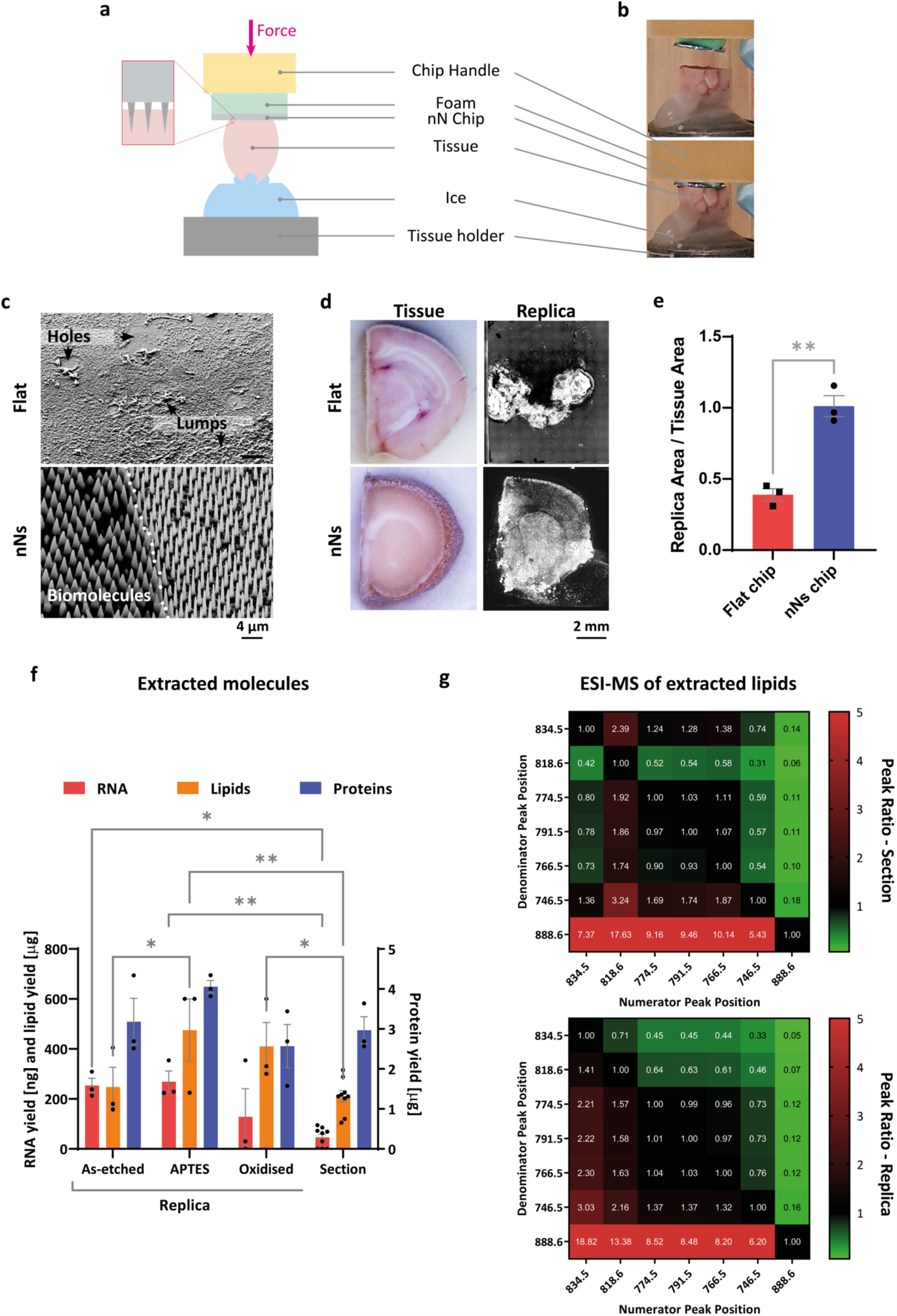
Generating a molecular replica using nanoneedles. (a) Schematics of the handling system, interfacing nanoneedles with tissue. (b) Pictures of the handling system approaching (top) and interfacing (bottom) nanoneedles with tissue. (c) Scanning electron microscopy images of flat and nanoneedle chips after interfacing with the tissue. The flat chip shows a non-uniform molecular adsorbate characterised by holes and lumps (top). The nanoneedle chip (bottom) shows the sharp edge between the uniform biomolecular adsorbate in interfaced area interfaced with the tissue on the left hand side, and the absence of molecular adsorbate outside of the area of interfacing on the right. (d) Photographic images of the sampled tissue and fluorescence microscopy images of the resulting molecular replica. The flat chips (top) show as partial replica which does not retain tissue morphology, while the nanoneedles (bottom) show a replica retaining the original morphology over the whole area of tissue interfacing. (e) Bar graph showing the ratio of the area of replicas over that of the original tissues. The data are shown as average with sem. Flat replicas have areas less than half of the tissue sampled. Nanoneedle replicas have areas comparable to the original tissues. (f) Bar graph comparing the amounts of RNA, lipids and proteins eluted from nanoneedles and tissue sections. Comparison across replicas of mouse brains generated on nanoneedles with different surface chemistries (as-etched, functionalised with APTES and oxidised) and a 10 µm section from the same mouse brain. The data are shown as average with sem. (g) Heatmaps showing the relative intensities from ESI-MS of lipid extract from section (top) and replica (bottom), for the seven most intense peaks. The relative intensities are overall preserved in the replica respect to the section.

We then assessed replica quality by comparing its molecular composition with that of the original tissue. For comparison we obtained a *reference tissue section* of 10μm thickness, proximal to the area of interfacing with nanoneedles. We evaluated the harvesting efficiency for RNA, proteins and lipids; three key classes of biomolecules for the study of living processes (Figure 2f). We compared replicas generated from nanoneedles as-etched, O_2_-plasma oxidised, and (3-aminopropyl)triethoxysilane (APTES) silanized to assess the role of surface polarity and charge on biomolecular harvesting efficiency^15,16^. Oxidation and APTES respectively generate negative and positive charges providing more polar surfaces than the as-etched porous silicon. For all the functionalizations considered, the amount of harvested biomolecules was detectable with conventional assays and compared favourably with the amounts that could be extracted from the reference tissue section. RNA harvesting was in the range of 210-350 ng while protein harvesting was in the range of 2.5-4.3 μg. Lipid harvesting was more efficient on APTES and oxidised nanoneedles ranging between 200-600 μg with respect to as-etched ones. This data indicated that plasma oxidized nanoneedles were suitable for our spatial lipidomic workflow, since their lipid harvesting efficiency compared favourably with a 10 μm tissue section, known to be imageable by DESI-MS.

Indeed, the lipidomic profile eluted from an oxidised nanoneedle replica was comparable to that of the reference tissue, as shown by the congruent relative intensities of most characteristic lipid peaks of brain tissue (Figure 2g) ^17,18^. Of note, only the putative phosphatidylserine (PS) peak at m/z 834.5 was preponderantly more abundant on nanoneedles. These data indicated that nanoneedles can provide a reliable replica of the lipidomic profile of a tissue.

### Lipid analysis on nanoneedles

We then assessed the suitability of nanoneedles for lipidomic imaging. DESI-MS imaging of porcine brain lipid extract deposited at decreasing concentrations onto nanoneedles and flat chips revealed comparable sensitivity across lipid species, as calculated from the limit of detection (LOD) for the most intense lipid peaks (Figure 3a). Putative identification assigned the peaks at m/z 788.5 and m/z 834.5 to PS, the peak at m/z 700.5 to phosphatidylcholine (PC), the peak at m/z 726.5 to phosphatidylethanolamine (PE), the peak at m/z 885.5 to phosphatidylinositol (PI), and the peak at m/z 888.6 to sulfatide (ST). The LODs for these species were within the same order of magnitude on the two substrates, ranging between 1 and 40 ng/cm^2^ on nanoneedles, and 0.3 and 15 ng/cm^2^ on flat surfaces, in agreement with previous DESI analysis^19^ (Figure 3b). The relative intensity of these characteristic peaks was preserved on nanoneedles with respect to those observed on flat substrates (Figure 3c).

**Figure 3:**
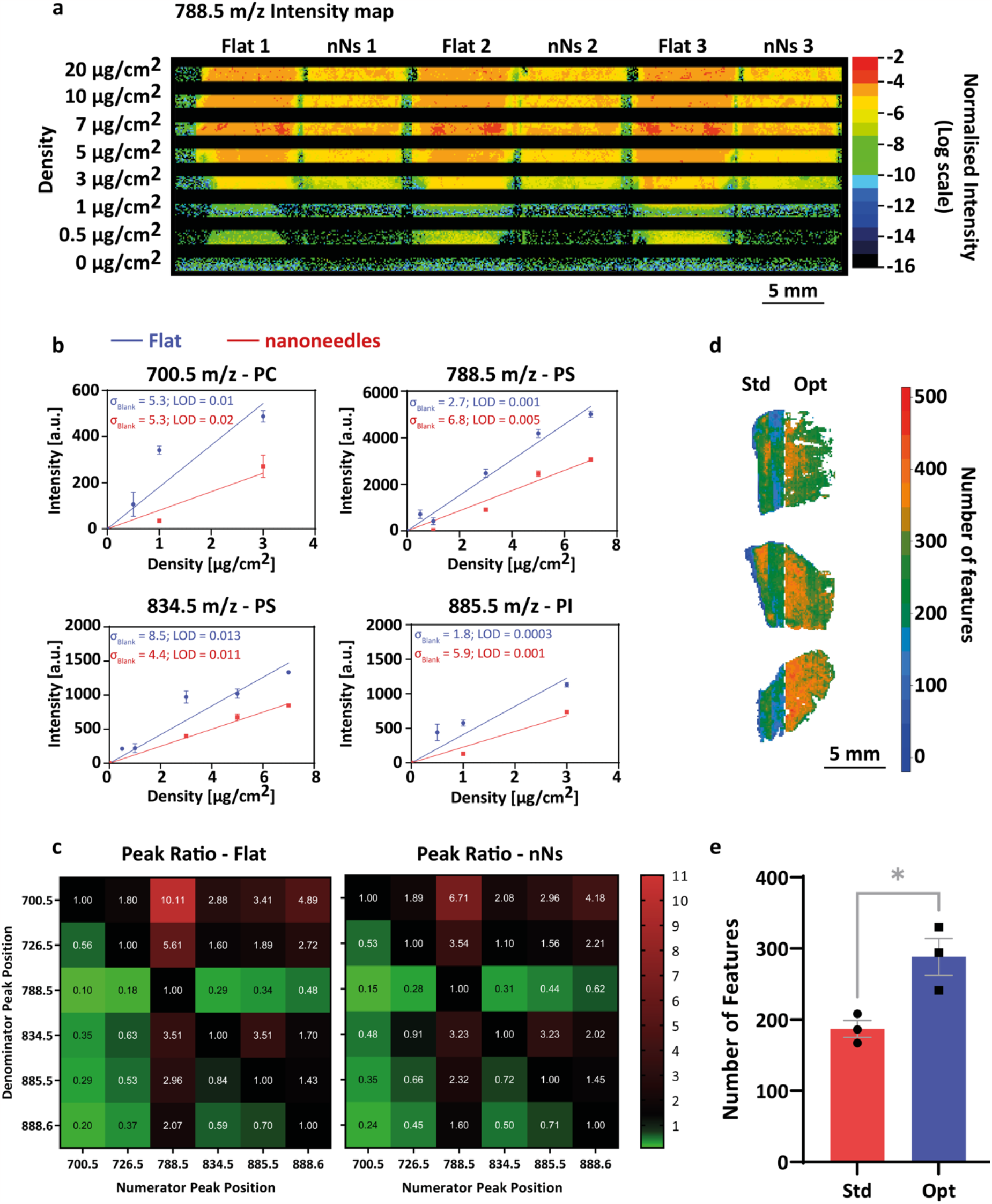
Lipid imaging on nanoneedles. (a) DESI-MS intensity maps for the highest intensity lipid-region peak at 788.5 m\z for porcine brain homogenate drop-deposited onto three flat and three nanoneedle chips, in the range of densities 0-20 μg/cm^2^. (b) Calibration curves calculated from the porcine brain homogenate DESI-MS imaging shown in panel (a), for representative lipid peaks at 700.5 m\z, 788.5 m\z, 834.5 m\z, and 885.5 m\z. Standard deviation of the blank (σ_blank_) and the calculated limit of detection (LOD) are reported, and fall within the same order of magnitude for flat and nanoneedles. (c) Heatmaps from flat and nanoneedle spectra at 7 µg\cm^2^, showing the relative intensities for the six most intense peaks in the lipid region. The relative peak intensity is overall preserved onto nanoneedles with respect to flat. (d,e) Optimization of DESI imaging conditions for nanoneedles: (d) Maps of the number of features from DESI-MS imaging of 3 liver replicas onto nanoneedles, imaged half with standard (std, left) and half with optimized (opt, right) DESI parameters. (e) Bargraph presenting as average with sem the quantification of the number of features before (std) and after (opt) the optimization of the DESI parameters, showing an increase from 180 to 290.

We optimised DESI-MS imaging conditions for tissue replica by combinatorial sweeping of key parameters to maximise the total number of detectable features in a replica of liver tissue (Figure S2)^20^. The starting values for the optimization parameters were standard settings for imaging of tissue sections, in agreement with the literature. The optimised settings were tested by imaging three replicas of mouse liver tissue. In each sample, half of the replica was imaged using standard settings, while the other half used optimised settings (Figure 3d). The parameter optimization yielded an increase of 53% in the total number of detectable features in the lipid spectral region (Figure 3e).

This data indicated that nanoneedles are a suitable substrate for lipidomic analysis with comparable performance to a conventional flat substrate for the detection of metabolites following DESI-MS.

### Spatial Lipidomics of murine glioma replicas

We assessed the quality of lipidomic imaging on replicas by comparing the spatial distribution of lipids of a murine brain replica and its proximal tissue section. Based on established DESI-MSI analytical workflows, the 1000 peaks with top intensity in the lipid region were selected for the analysis^21,22^. Noise removal was used to filter out peaks with high frequency and low intensity. Peaks within m/z 0.02 across different samples were matched.

We compared the ability of replica and reference section to discriminate tissue morphology and composition^14^. Hierarchical Cluster Analysis (HCA) revealed that the top two clusters of both section and replica appear to overlap with the areas of grey and white matter within the mouse brain architecture (Figure 4a). The cluster labelled as “white matter” overlapped with brain regions composed by highly myelinated fibers, matching the patterns of expressions of myelin genes such as myelin basic protein MBP in areas like the corpus callosum, while the cluster labelled “grey matter” seemed to map onto areas with higher cell bodies densities with less myelinated fibers like the cortex^13^. The average lipid composition of the two clusters confirmed this morphological identification (Figure 4b,c). Comparing relative lipid abundance identified species characteristic of white and grey matter respectively. The higher abundance of the putative sulfatide ion at m/z 888.6 and lower abundance for the putative PS peak at m/z 834.5 in white matter well agrees with established DESI-MS brain biomarkers^17^. Mapping the relative intensity of these two peaks across the sample accurately outlined white and grey matter in the replica as well as in the section (Figure 4d). The congruent distribution of these lipid species across replica and section established the feasibility of generating a lipidomic profile map of a nanoneedle replica of mouse brain.

**Figure 4:**
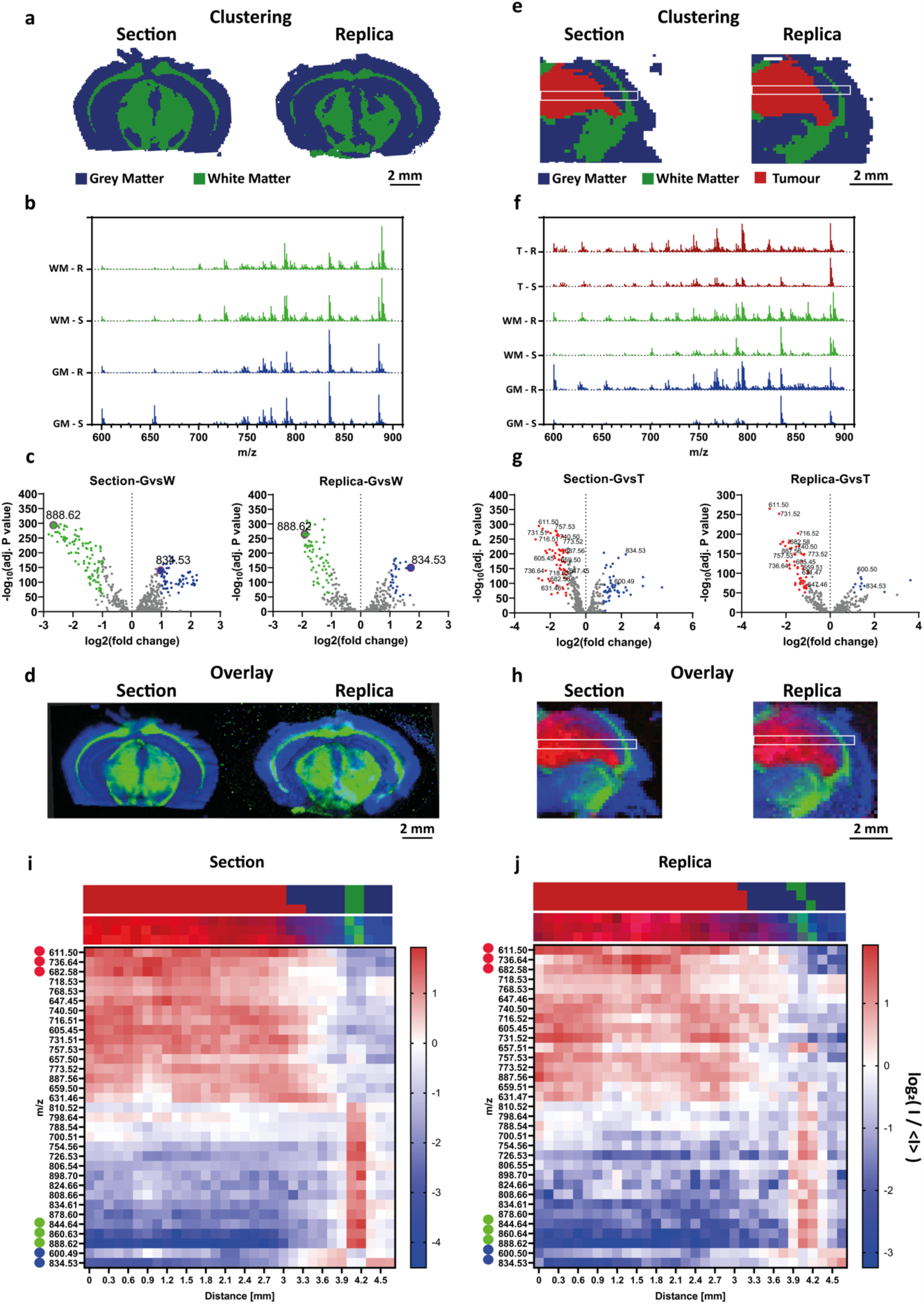
Spatial lipidomics of murine glioma replicas. (a) Maps of the two top clusters, obtained by hierarchical cluster analysis (HCA) of the DESI-MSI dataset of a murine brain section and its replica. The maps correlating with the morphological structure of grey (blue cluster) and white matter (green cluster). (b) Average spectra showing normalized counts vs m\z, for the top two clusters associated to grey (GM) and white matter (WM) in section (S) and replica (R). (c) Volcano plots showing the differential lipid distribution in grey matter vs white matter. The plots indicate a preserved key role for the established phosphatidylserine ion at 834.5 m\z and sulfatide ion at 888.6 m\z as biomarkers of grey and white matter across both section and replica. (d) Map overlay of the total ion count-normalized intensity for the peaks at 834.5 m\z (blue) and 888.6 m\z (green), showing the characteristic morphology of grey and white matter, preserved in the replica and the section. (e) Maps of the top three clusters, obtained by HCA of the DESI-MSI dataset of a tumour-bearing murine brain section and replica, mapping to GM (blue cluster), WM (green cluster), and tumour (red cluster). The white boxes show the regions considered for the analysis of the lipid abundance across the three tissues. (f) Average spectra showing normalized counts vs m\z, for the 3 clusters associated to GM, WM, and tumour (T) in section (S) and replica (R). (g) Volcano plots showing the differential lipid distribution in grey matter vs tumour, with highlighted the 16 (part of the 33 found by considering also white matter) most representative lipid peaks in both section and replica. (h) Map overlay of the cumulative total ion count-normalized intensity for the peaks at 600.5 and 834.5 m\z (blue), 844.6, 860.6, and 888.6 m\z (green), and 611.5, 736.6, and 682.6 m\z (red) discriminating grey, white matter, and tumour in section and replica. The white boxes show the regions considered for the analysis of lipid abundance across GM,WM, and T. (i-j) Heatmaps showing the relative abundance along the major axis of the white box shown in panels (e,h) of the 33 lipid peaks identified by differential analysis (g) within the section (i) and the replica (j). The distribution of these lipids well defines GM,WM, and T in both section and replica. The two strips over each heatmap show the cluster assignment (top) and relative lipid intensity (bottom) for the corresponding set of pixels (e,h).

We then assessed the replica’s ability to characterise a cancerous lesion within a murine glioma model. As observed in DESI-MS imaging of sections from tumour bearing mouse brain ^14^, in our sample the top three HCA clusters classified white matter, grey matter and tumour with comparable effectiveness in section and replica (Figure 4e). When analysing a dataset including all spectra across tissue and replica, HCA similarly discriminated tumour, white and grey matter, highlighting the correspondence of the lipid profile on replica and section. The assessment of inter-cluster variance by principal component analysis confirmed that replica and section present congruent characteristic features, and that inter-cluster variance can map tumour margins and transitions between tissues with analogous efficacy in the replica and the section (Supplementary Information, Figure S3). Analysis of the average mass spectra for each cluster enabled comparing the lipid composition of white matter, grey matter and tumour and identifying the most relevant among differentially abundant species (Figure 4f,g, Figure S4). In agreement with the DESI-MSI literature, both section and replica presented clusters associated to white matter with abundant peak at m/z 888.6, putatively assigned to sulfatide, while the grey matter clusters showed a prominent peak at m/z 834.5 putatively assigned to PS. The tumour cluster showed an increase in the abundance of m/z 611.5 (putatively cholesterol esters, CE) and m/z 716.5 (putatively PS/PE). The intensity map for the characteristic species identified through the volcano plots effectively discriminated tumour, white and grey matter in the replica as well as in the section (Figure 4h), indicating the effectiveness of the replica in providing a lipidomic map of the original tissue in presence of a cancerous lesion. To test the reproducibility of our results we obtained lipidomic maps of replicas and reference sections for two additional, distinct tumour-bearing animals, showing an analogous ability to correctly classify tumour, white, and grey matter by hierarchical cluster analysis (Figure S5).

We compared the spatial distribution of analytes in the replica and the proximal section by evaluating lipid abundance along a line spanning the length of the sample, crossing white matter, grey matter and tumour, corresponding to the major axis (length) of the white box in Figure 4h. The relative abundance of a panel of the 33 most representative lipid species, as highlighted by the volcano plots, was calculated for every pixel along the length of the white box, and averaged over the three pixels of its width. The heatmap from the section showed a clear pattern of relative abundance for the tissue-associated clusters of lipids when transitioning between tumour, grey and white matter (Figure 4i,j). The first transition, between tumour and grey matter, occurred at 3.1 mm from the left edge. The tumour was rich in lipids at m/z 611.5 (putatively cholesterol esters, CE), m/z 736.6 and m/z 682.6 (putatively ceramides, Cer), m/z 731.5 (putatively sphingomyelin, SM) and m/z 740.5 (putatively PS / PE). Lipids at m/z 600.5 (putatively ceramides) and m/z 834.5 (putatively PS) were abundant in the grey matter. At 4 mm the sample presented a 0.3 mm layer of white matter abundant in lipids at m/z 844.64 (putatively PS/PE), at m/z 860.63 (putatively PS) and m/z 888.62 (putatively sulfatide) followed by a grey matter region stretching from 4.3 mm to 4.7 mm, with the same pattern of abundance as the other grey matter region. The heatmap from the replica showed the same spatial pattern and lipid abundance of the section (Figure 4j). In the replica, the first transition between grey matter and tumour arose at 3.1 mm, followed by a 0.3 mm layer of white matter at 4 mm, before returning to grey matter between 4.3 mm and 4.7 mm. The same agreement in spatial distribution of relative lipid abundance across replica and section arose when comparing other regions within this sample, including the corpus callosum and the peduncle region of the white matter feature (Figure S6).

These data indicate that a nanoneedle molecular replica of a tumour-bearing mouse brain can map the biomolecular architecture of the original tissue. The quality of information provided by the replica enables unsupervised classification of individual pixels with comparable fidelity to the original sample. This information enables evaluating the relative abundance of metabolites and mapping their distribution with comparable quality to that attainable from the original sample.

### Spatial Lipidomics of human glioma replica

Having validated the molecular replica approach to map and discriminate the lipid composition of brain with comparable quality to the analysis of tissue sections, we applied this approach to the analysis of a human glioma biopsy. The histopathological analysis of the lesion showed three distinct regions classified as healthy brain infiltrated by tumour (I), bulk tumour (T), and necrosis (N) (Figure 5a). HCA could distinguish these three regions in the replica as well as in the reference section (Figure 5b,c). From the unsupervised classification it was possible to identify the characteristic lipid composition of each region for both samples (Figure 5d). This analysis enabled identifying individual species characteristics of the three regions for both replica and section. The peak putatively associated to ceramides at m/z 600.5 characterised the infiltrated tumour, while sulfatide at m/z 888.6 distinguished the bulk tumour and phosphatidylcholine at m/z 680.5 preferentially associated with areas of necrosis (Figure 5e). Mapping the distribution of the characteristic peaks across the section and the replica outlined the three regions identified by histology. From the volcano plots we further identified 10 lipids, whose change in abundance correlated with the transition between regions, both in section and replica (Figure 5f).

**Figure 5:**
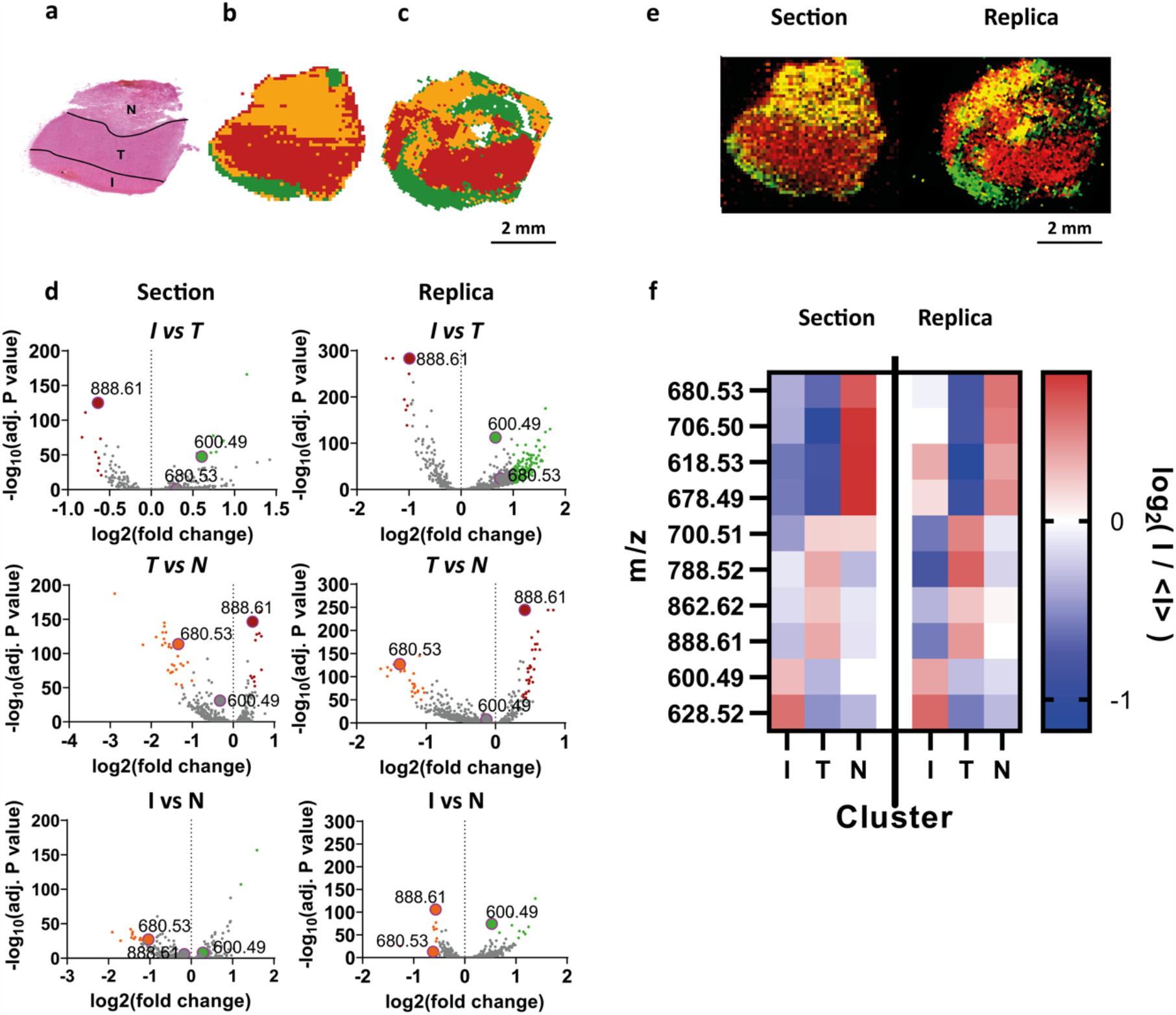
Spatial lipidomics of human glioma replica. (a) Bright-field image of H&E staining of human biopsy section from glioma patient, presenting three regions identified as infiltrated tumour (I), core tumour (T), and necrosis (N). (b, c) Maps of the top three clusters, obtained by HCA of DESI-MS images of section (b) and replica (c), showing good association with the regions displayed by the H&E. (d) Volcano plots showing the differential analysis of lipid abundance in I, T and N. Both section and replica indicate a key role for ceramide ion at 600.5 m\z (I), sulfatide ion at 888.6 m\z (T), and phosphatidylcholine ion at 680.5 m\z (N) at discriminating the main regions within the sample. (e) Map overlay of the total ion count-normalized intensity for the peaks at 600.5 m\z (green), 888.6 m\z (red), and 680.5 m\z (yellow). (f) Heatmap showing the relative intensity of the 10 most representative lipid species identified by the differential abundance analysis, that define the 3 tissues (I, T, and N) both in section and replica.

These data indicates that DESI-MSI of molecular replicas of human glioma can provide a reliable spatial lipidomic map of the original tissue of interest.

### Human glioma replicas are proxies of the tissue’s molecular profiles

To systematically assess whether the molecular profiles of tissue replicas can reliably approximate those of their reference tissue sections we collected a patient dataset consisting of DESI MSI maps of a replica and its reference tissue section from 23 human glioma tumours with known prognosis (Methods, Table S1). The dataset included one additional section from each of samples HG 11 and HG 12 alongside one additional replica from each of samples HG 6 and HG 18, for a total of 25 sections and 25 replicas. We initially looked at the distribution of m/z 600.5, m/z 888.6 and m/z 680.5. (Figure 6a) These peaks were chosen as they respectively associated to infiltrated, bulk and necrotic tumour in our analysis of sample HG 25 (Figure 5e). Their distribution across a representative selection of samples identified distinct regions within the lesions for both section and replica.

**Figure 6:**
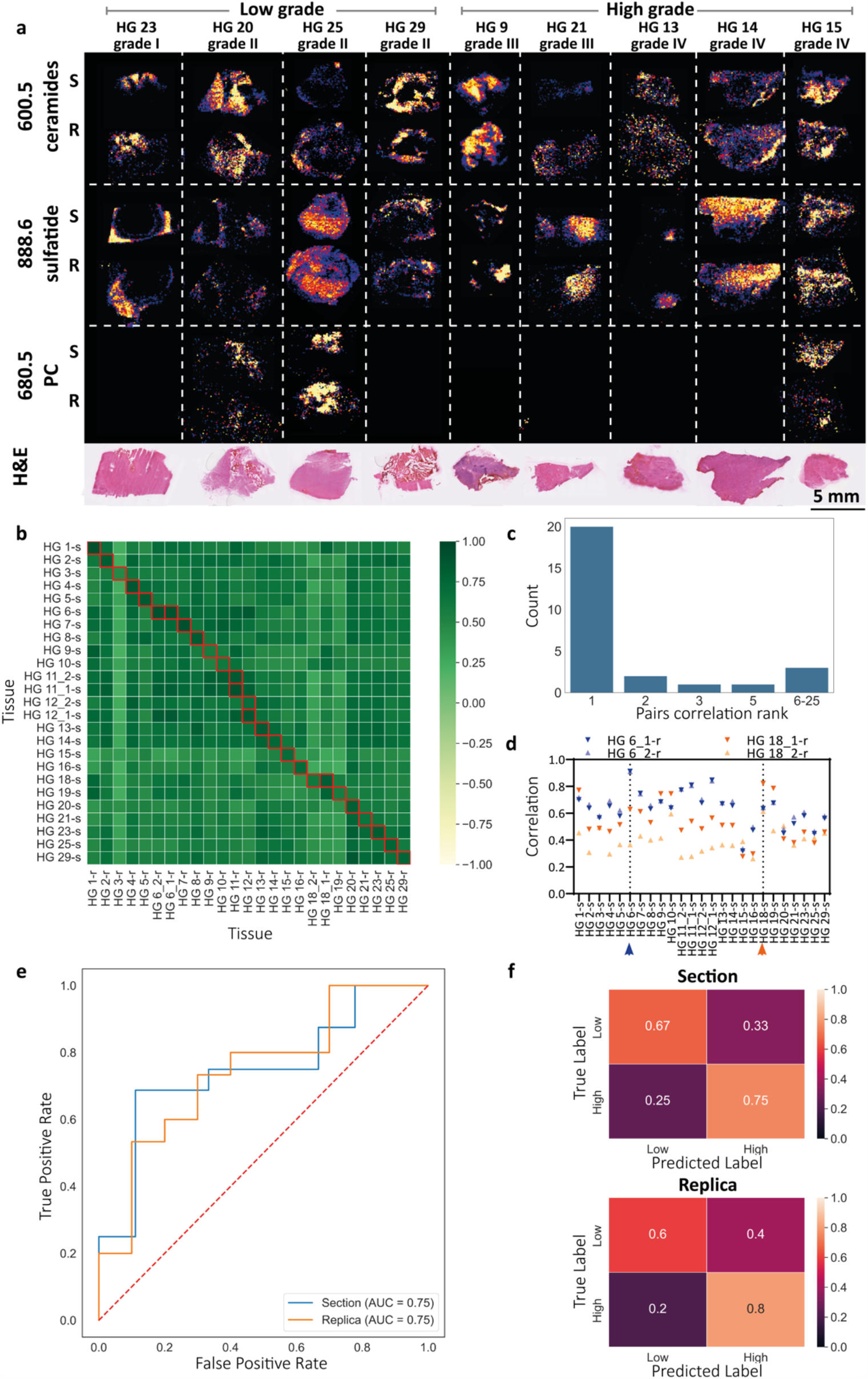
Grading of human gliomas using molecular replicas. (a) Map of the total ion count-normalized intensity for a selection of human glioma replicas (R) and reference sections (S). The images display the ceramide peak at m\z 600.5, sulfatide peak at m\z 888.6, and phosphatidylcholine peak at m\z 680.5. The respective sample morphology is shown in the H&E section at the bottom. (b-d) Similarity between tissue reference section and replica. (b) Heatmap of correlation values between all 25 tissue sections and 25 replicas of 23 biopsy samples. Individual biopsy samples are labelled HG followed by a sequential number. Multiple sections or replicas from the same biopsy are labelled with a trailing underscore and sequential number. Sections are denoted with a trailing s and replicas with a trailing r. The red boxes represent matched tissue sections and nanobiopsy replicas. (c) Count plot of correlation rank between matching tissue sections and nanobiopsy replicas. Most replicas are the first correlated sample to their section, and the overall distribution greatly differs from a random correlation. (d) Graph showing the Pearson correlation of the the nanobiopsy replicas HG6_1-r, HG6_2-r, HG18_1-r, HG18_2-r, with all 25 tissue sections. The blue arrow highlights the location of the reference HG6-s tissue section and the orange arrow highlights the location of the reference HG18-s tissue section. Correlation between multiple replicas and their matched section was higher than any other sections. (e,f) Performance of a deep neural network-based binary classifier for glioma grade applied to tissue sections and nanobiopsy replicas. (e) Receiver operator characteristics curves for the tissue sections (blue) and replicas (orange) display an area under the curve for the receiver operating characteristic (AUC-ROC) of 0.75 and 0.75 respectively. (f) Normalised confusion matrices for tissue sections and nanobiopsy replicas display comparable predictive power. Upper confusion matrix for sections. Lower confusion matrix for replicas.

To overcome inconsistencies in spectra dimensions and noise we developed a preprocessing pipeline consisting of tumour segmentation, spectra correction in respect to the non-tissue regions, normalisation and quantization (Methods, Figure S7). After preprocessing, all spectra shared a common and meaningful representation suitable for direct comparison and evaluation. To assess whether the molecular profile of a replica accurately represented its reference section, we performed a correlation-based analysis. For all possible section-replica pairings, we evaluated the Pearson correlation between their mean spectra. This analysis produced a matrix of size 25×25 where bin (i,j) indicated the correlation between section i and replica j (Figure 6b). In general, a replica and its reference tissue section showed high correlations, indicating that the characteristic molecular profile of glioma biopsies was maintained in the replicas across the DESI-MS spectrum. To quantitatively assess whether a replica was most similar to its matched section we calculated the correlation ranking order. For each replica, we ranked all 25 sections according to their correlation. Over the 27 matched pairs indicated by the red boxes in figure 6b, the probability of identifying the true matched section was 0.74, and that of being in the top five correlative sections was 0.89, much higher than the expected random probability of 0.04 and 0.2 respectively (Figure 6c). To determine whether multiple replicas from the same section maintained their molecular signature, we compared the correlation between the two replicas of the same tissue (HG 6 r1, HG 6 r2 and HG 18 r1, HG 18 r2) with their reference section against all other sections. The correlation of each of the replicas to their reference section was higher than any of the other sections (Figure 6d). These results suggest that multiple replicas from the same section capture a similar and representative molecular profile of the corresponding tissue section.

### Machine learning inference of human glioma grade

We next asked whether the molecular profiles of tissue sections can predict the disease state, and if so, whether their replicas can match such predictive value. Based on their histopathological annotations, we partitioned each glioma biopsy to either high (grade III-IV, N = 15 replicas, 16 sections) or low (grade I-II, N = 10 replicas, 9 sections). We trained deep neural networks (methods) on the single spectra from sections/replicas, to discriminate between high and low grade glioma. This approach of single spectra classification was taken, because of the limited cohort size and under the assumption that the grade was homogeneous within each biopsy. Because of the limited size of our cohort, and to avoid data contamination of having spectra from the same patient or imaging session (batch) in both training and testing, we assessed the tumour grade classification performance using leave-one-batch-and-patient-out analysis (Table S2). This included multiple rounds of training and testing, where each round with the sections/replicas imaged in one session designated as the test set, while the rest of the data were used as the training and validation sets (Methods). The classification performance of our model indicated that glioma grade could be effectively predicted from the molecular profiles of tumour sections and their replicas (Figure 6e). The area under the curve (AUC) for the receiver operating characteristic (ROC) was comparable (0.75 Section, 0.75 Replica) as were the balanced accuracy (0.71 Section, 0.70 Replica) and the confusion matrices (Figure 6f, Table S3). The section and replica models agreed on 21/27 section-replica pairs further suggesting that the models captured similar discriminative information in section and replica spectras (Table S1). Testing the model trained on the sections using the replica dataset, still yielded good performance metrics with AUC 0.71 and balanced accuracy 0.60, further supporting the similarity between the molecular profile of replicas and their reference tissues (Table S3). These results established that the molecular replica generated using nanoneedles can replace a tissue section for disease state classification.

## Discussion

We developed a workflow that uses a nanoneedle chip to generate a reliable molecular replica of a tissue, and to map its lipid composition by DESI-MSI for glioma characterisation and classification. The molecular replica preserved the relative lipid composition of the original samples and their distribution. When generating a replica of a mouse brain bearing an orthotopic glioma, the molecular information within the replica was sufficient to map white matter, grey matter and tumour with accuracy comparable to a reference tissue section. Further analysis indicated that the replica and its reference section shared key differentially abundant lipids between white matter, grey matter and tumour, which were preserved on a pixel-by-pixel basis. When applied to human glioma biopsies, DESI-MSI analysis of a replica could map the infiltrated, bulk and necrotic regions of a tumour, enabling to identify with displaying a comparable differential lipid abundance to the reference section within these three regions. The analysis of 23 human gliomas showed that the lipid composition of replicas was highly correlated to that of their reference sections. The replicas could predict disease grade using a deep neural network on a pixel-by-pixel assessment, in both sections and replicas.

These findings indicate that nanoneedles can serve as a platform for tissue-wide, minimally invasive molecular diagnostics. In particular the ability of molecular replicas to identify biomarkers for tissue regions and lesion state is comparable to a tissue section. The spatial distribution of the biomarkers is well preserved on the replica, serving to accurately outline lesions, their state and define margins. Furthermore, the molecular information provided by the replica can effectively fingerprint tissue state, provide sufficient data for unbiased classification that performs on par with the original tissue. Since nanoneedle application to tissues is non-perturbing^23,24^, these results indicate the way to the noninvasive collection of patient and *in-vivo* samples for spatial molecular diagnostics.

Moving forward, by accessing a larger, and more tailored set of human glioma samples it should become possible to provide a more granular stratification of samples by discriminating glioma subtypes and known mutations such as astrocytic or oligodendocytic nature of the malignancy and isocitrate dehydrogenase (IDH) involvement, which are detectable using DESI-MSI^25^. Increasing the sampling resolution of imaging on replicas and faster imaging modalities should also be explored as the capability of spatial lipidomics consistently improves.

## Methods

### Fabrication of nanoneedles

Nanoneedles were fabricated according to our established protocols ^23,26^. The first step was photolithography patterning of 4’’ 0.01-0.02 Ω·cm, B-doped Si wafer (University Wafers Inc, USA) coated with 120-160 nm of epitaxial Si-rich Si_3_N_4_ (University of Edinburgh). Following dehydration in oven at 200 °C for 20 min, the photoresist NR9-250P (Futurrex, USA) was spin-coated following 3 spinning steps: 5 s / to 500 rpm / 1000 rpm/s; 40 s / at 4000 rpm / 5000 rpm/s; 0.1 / to 0 rpm / -5000 rpm/s. After spin-coating, the wafer was pre-baked on hotplate at 70 °C for 180 s. A square lattice pattern of 0.6 µm dots spaced 2 µm was transferred to the photoresist, by 2.9 s exposure with a mask aligner (K-SUSS MA6). After exposure, the wafer was post-baked on hotplate at 100 °C for 60 s. The photoresist was developed for 12 s in 3:1 RD6:H_2_O, immersed in water to stop the development, rinsed with excess water and dried under N_2_ flow. The second step was the pattern transfer from the photoresist to the Si_3_N_4_ by CHF_3_ plasma RIE, (Oxford Instruments, NGP80), with 50 sccm CHF3, 5 sccm O2, 150 W forward power, 55 mTorr pressure, for 150 s. The photoresist was stripped with acetone, rinsed with IPA and N_2_ dried. The wafer was dipped in 100 mL of cleaning solution 10 %v/v HF (Honeywell, 95299) for 2 min. The third step was the electroless deposition of Ag nanoparticle for 2 min, from 20 mM AgNO_3_ (AgNO3, Sigma-Aldrich 99.9999%) dissolved in 100 mL 10 %v/v HF. Then, DI water and IPA rinsing, N_2_ drying. The fourth step was MACE, for porous Si pillars fabrication, by dipping the wafer in 400 mL of etching solution H_2_O_2_:HF(10%v/v) 1:99 for 7.5 min. To stop the etching, the wafer was immersed in excess DI water, then rinsed with DI water and IPA, and dried under N_2_ flow. The fifth step was Ag removal, by dipping in gold etchant solution (Aldrich, 651818) for 10 min. DI water and IPA rinsing, N_2_ drying. The last step was the nanoneedle shaping, by SF_6_ plasma RIE (Oxford Instruments, NGP80), with 20 sccm SF_6_, 300 W, 100 mTorr, for 120s. After fabrication, the wafer was diced in 0.8 × 0.8 cm^2^ chips (Disco, DAD3220).

### Animal tissues

Procedures were carried out under compliance of the Animals (Scientific Procedures) Act 1986, Home Office license and KCL ethical review approval. The murine brain and liver tissue used for interfacing strategy definition, molecule harvest total quantification and DESI-MS imaging optimisation, were collected from wild type, adult CD1 mice, when the culling of the animal, to collect other organs, was already planned from researchers in the Centre for Craniofacial and Regenerative Biology (KCL, London, UK). The brain tumour murine model consisted in C57BL/6 J ^27^, injected with neural stem cells with concomitant Nf1, Pten, EGFR-vIII mutations (NPE) ^28^. In brief, mice ∼ 6–8 weeks age were injected with 400k cells into the right hemisphere. Starting from the bregma the stereotactic coordinates were as follows: 3 mm anterior, 1mm lateral, and 3 mm deep^29^. Roughly 2–3 weeks after the implantation of the tumours, animals were humanely sacrificed and the brains were collected and frozen in - 20° C. All animal experiments were performed under the authority of project licence (PBE6EB195 - Prof. Al Jamal Khuloud) and personal licence (I13B24EFC – Miss. Nadia Rouatbi) granted by the UK home office and the UK CCCR Guidelines (1998) in compliance with the UK home office (1989) code of practice and EU directive 2010/EU/63 for the housing and care of Animals used in Scientific Procedures.

### Human Biopsies

Human Tissue samples from patients with ragiological evidence of a brain tumour were obtained from consented patients undergoing surgery at the Department of Clinical Neurosciences, NHS Lothian, Edinburgh. Tissue sample collection was approved by a local regional ethics committee (Lothian NRS Bioresource 20/ES/0061). Samples were snap frozen and stored until analysis at -80 °C. Tumour diagnosis was established from histological analysis.

### Cryosectioning of tissue

The tissue sections were collected with a Bright OTF5000 cryotome. The chamber and specimen temperature were set at -20 °C. The sections were collected onto glass slides, superfrost® plus slides (fisher scientific 12625336).

### Generation of tissue replicas onto nanoneedles

The specimen was transferred outside the cryostat and the chip was placed in contact with the flat surface for less than 1 s, to allow the harvesting of biomolecules onto nNs. After the replica generation, the nanoneedles were left to dry in air 1-2 min and transferred onto the quick freeze bar inside the cryostat until storage in -80 °C.

### Lipid staining of tissue replicas on flat and nanoneedles

Lipophilic carbocyanine dye Neuro-DiO (Biotium, 30021) was used to stain the lipid of tissue replicas according to the manufacturer protocol. In brief, the tissue replicas were first fixed using paraformaldehyde (4% in PBS) for 10 minutes at room temperature. After PBS (1x) washing 3 times, enough freshly prepared Neudo-DiO (in 1x PBS) staining solution was added to cover the nanoneedles, then incubated 30 minutes in the dark at room temperature, and finally washed 3 times with PBS. Tile scan images were acquired by Leica DMi8 inverted microscope (Leica Microsystems GmbH) with 20x 0.4 NA air objective.

### Histology of tissue sections

Tissue sections were processed by H&E using an automated LeicaBOND tissue processor. After completion of the mounting at 60°C overnight, the stained sections were imaged by the slide scanner Hamamatsu Nanozoomer. The software NDP view 2 (Hamamatsu) was used for visualization and annotations of H&E stained slides.

### Extraction and total quantification of biomolecules

#### RNA

All surfaces were wiped with RNase AWAY™ (Thermo Scientific™ 10666421). Biomolecules were extracted with 300 µL TRIzol. To assist the elution of biomolecules from 1 cm^2^ nNs and glass chips, ceramic beads (MP Biomedicals, Lysing matrix D, 2 mL tube, 6913050) with 20 s, 4 m/s tissue homogenisation (MP Biomedicals FastPrep-24) were used. The solution was centrifuged at 16000 x g for 1 min and the supernatant was collected, to remove the nNs and glass residue. Direct-zol™ RNA MiniPrep Plus (Zymo research, R2070S) was employed for RNA extraction. An equal volume of ethanol 100% was added to the sample before transferring it to a Zymo-Spin™ IIICG Column (Zymo research, C1006). The sample was centrifuged at 16000 x g for 30 s, and the flowthrough underwent to protein extraction. The column was transferred to another RNase free tube and underwent 2 pre-washes and 1 wash step using the buffers provided with the columns, as per manufacturer instruction. Finally, the RNA was collected in RNase free water. Qubit™ RNA HS assay (Thermo Fisher Scientific, Q32852) was employed for total RNA detection, following the user manual from the manufacturer, using a Qubit 3.0 fluorometer (Invitrogen™ Q33216).

#### Proteins

The flowthrough from the RNA extraction in the column was incubated for 30 min on ice, after adding 4 volumes of cold acetone (-20 °C). After centrifugation at 20,000 rcf for 10 min, the protein pellet underwent ethanol wash and centrifugation at 20,000 rcf for 1 min. The protein pellet was air dried for 10 min at room temperature, then resuspended in 100 µL protein storage buffer: 4 M urea (Sigma-Aldrich, U5128), 1 % SDS (Sigma-Aldrich, L3771). Proteins were kept at -80 °C until quantification. The detergent was removed from the samples in storage buffer by loading 100 µL of sample in pre-dispensed HiPPR™ spin columns (Thermo Scientific™, 88305), then following the user manual from the manufacturer. Desalting was performed by loading 100 µL of sample in Pierce™ Protein Concentrators PES, 3K MWCO (Thermo Scientific™, 88512), following the user manual from the manufacturer. The protein concentrator allowed to change the storage buffer to 100 µL of 0.1 M sodium borate (Sigma-Aldrich, HT1002), pH 9.3, for total quantification by CBQCA protein quantitation kit (Invitrogen™, C6667). A standard curve using bovine serum albumin (Sigma, A9647) was built following the manufacturer instruction. Samples and standards were processed according to the user manual from the manufacturer. Fluorescence was measured using the CLARIOstar® Plus platereader (BGM Labtech), with excitation waveband 465 ± 15 nm and emission waveband 550 ± 20 nm.

#### Lipids

Biomolecules were extracted with 200 µL CH_2_OH-H_2_O mixture or CHCl_3_. To assist the elution of biomolecules from nNs and glass chips ultrasonication or ceramic beads (MP Biomedicals, Lysing matrix D, 2 mL tube, 6913050) with 20 s, 4 m/s tissue homogenisation (MP Biomedicals FastPrep-24) were used. Samples were immediately placed on ice. The solution was centrifuged at 16000 x g for 1 min and the supernatant was collected, to remove the nNs and glass residue. The ratio of CHCl_3_-CH_2_OH-H_2_O was brought to 1:2:0.8 and the sample mixed, then CHCl_3_ was added to obtain ratio 2:2:0.8, vortexed for 30 s and allow phase separation. The aqueous layer was removed. The organic phase was transferred into new tubes for quantification by Nile Red staining. A stock solution was prepared for Nile Red (NR) (9-diethylamino-5H-benzo[α]phenoxazine-5-one, C_20_H_18_N_2_O_2,_ Sigma Aldrich 72485) in acetone 1:40 v/v. The samples were dried under N_2_ flow to 1/10 of the original volume or until complete evaporation. A volume of 200 µL of water was added to the samples. Immediately after vortexing the samples, 2 µL of NR stock solution was added to the samples. The samples were then vigorously vortexed for 1 min to obtain a microemulsion. The samples were finally transferred to a well plate for analysis. Fluorescence was measured using CLARIOstar® Plus microplate reader, with excitation waveband 530 ± 15 nm and emission waveband 612 ± 100 nm.

### ESI-MS analysis of lipid extracts

For the analysis of the lipids by ESI-MS, 200 µL of chloroform: methanol (2:1) were used to extract the lipids from brain replica and section. 70 µL of LC-MS grade water (Fisher Scientific, 10728098) were added to each sample and the sample was vortexed for 10 s, then centrifuged for 3 min at 3000 rpm at 4 °C to promote the phase separation. The aqueous layer was removed, and the organic layer transferred to new tubes. The sample was evaporated under N_2_ flow, then a loading buffer was added. To prepare the loading buffer, 4.625 mg of ammonium acetate (Sigma-Aldrich, 73594) were dissolved in 210 μL of isopropanol: acetonitrile: water (2:1:1). After adding the loading buffer, the samples were centrifuged a last time at 13000 rpm for 3 min, to remove possible tissue residues. LC-MS grade isopropanol (Fisher Scientific, 10684355) and acetonitrile (Fisher Scientific, A955-1) were used.

ESI-MS was performed using a Xevo® G2-XS Tof mass spectrometer (Waters™) with ESI ion source controlled by MassLynx 4.2 software. The instrument was operated in sensitivity mode and negative polarity. The spectra were acquired with 20,000 FWHM (mass accuracy ≤ 1 ppm) mass resolution, over the range m/z 50-1200. The cone temperature was set to 100 °C, the cone voltage was set to 40 V. The solvent composition was CH_2_OH:H_2_O 0:1, with 0.1% v/v formic acid. The samples in loading buffer isopropanol: acetonitrile: water (2:1:1) 10mM ammonium acetate were injected with flow rate set at 5 μL/min.

### Optimisation of DESI-MS Imaging on nanoneedles

The total lipid extract from porcine brain (Avanti® Polar Lipids, Inc. 131101C) was diluted in methanol to obtain concentrations between 8 and 320 ng/µL. For each concentration 10 µL were drop deposited onto the nNs and flat surface. The surfaces were dried in air. 3 replicas and 3 sections were imaged for each concentration. The spectra were acquired by a Xevo® G2-XS Tof mass spectrometer (Waters™, Milford, MA, USA) equipped with DESI ion source (Waters™, Milford, MA, USA) controlled by MassLynx 4.2 software. The instrument was operated in sensitivity mode and negative polarity. The spectra were acquired with 20,000 FWHM (mass accuracy ≤ 1 ppm) mass resolution, over the range m/z 50-1200. The cone temperature was set to 150 °C, the cone voltage was set to 50 V. HD Imaging 1.4 software (Waters™, Milford, MA, USA) was employed before image acquisition, to set up imaged region of interest, pixel size 100 µm and scan rate 100 µm/s.

After data collection HD Imaging was employed to select the 1000 most intense peaks across the spectral map, in the lipid region (m/z 600-900). HDI exported as .txt file the map of spectra with reduced number of peaks. MATLAB (MathWorks, Inc., Natick, MA, USA) was used to parse the data into a format accessible by the R package used for the analysis of the spectral maps.

For the analysis of the spectral maps, R environment was used to adapt functions from *hyperSpec* package ^30,31^. For each chip, average intensity values were calculated for selected peaks from spectral maps composed by 12 × 19 pixels. The reported value and standard deviation are relative to the mean of the average intensities for the 3 flat and 3 nN chips, respectively. The LODs were calculated as 3 standard deviations of the blank over the linear coefficient of the curve fitting the linear region of the calibration curve.

### DESI-MS Imaging

DESI-MSI was performed using a Xevo® G2-XS Tof mass spectrometer (Waters™, Milford, MA, USA) with DESI ion source (Waters™, Milford, MA, USA) controlled by MassLynx 4.2 software. The instrument was operated in sensitivity mode and negative polarity. The spectra were acquired with 20,000 FWHM (mass accuracy ≤ 1 ppm) mass resolution, over the range m/z 50-1200. The cone temperature was set to 150 °C, the cone voltage was set to 50 V. HD Imaging 1.4 software (Waters™, Milford, MA, USA) was employed before image acquisition, to set up imaged region of interest, pixel size (between 50-100 µm) and scan rate 100 µm/s.

After data collection HD Imaging was employed to select the 1000 most intense peaks across the spectral map, in the lipid region (m/z 600-900). The map of spectra with reduced number of peaks was exported as .txt file. MATLAB (MathWorks, Inc., Natick, MA, USA) was used to parse the data into a format accessible by the R package used for the analysis of the spectral maps.

For the analysis of the spectral maps, R environment was used to adapt functions from *hyperSpec* package ^30,31^. First, the spectra were normalised by total ion count (TIC). Second, the spectra associated with the background by HCA were removed. Third, the distribution of the peak intensities was generated from the average spectrum. Nonlinear least squares were employed to fit the distribution profile with the log-normal function and a threshold was defined for the denoising at 1σ from the mean. All the peak positions having intensity below the threshold in the average spectrum were removed. Third, merging of 2 datasets associated to section and replica. A matrix of the differences between peak positions (m/z) in the 2 datasets was defined. From the matrix of the differences, based on a defined tolerance of m/z 0.02, a logic matrix was defined by the criterium *difference < tolerance*. In case of multiple matches, the closest value was picked. The R package *hyperSpec* was employed to perform HCA and PCA. The graphics were prepared using GraphPad Prism 9 (GraphPad Inc., La Jolla, CA, USA). To plot the volcano plots, differences between 2 groups were calculated by student’s t-test. Peaks with log_10_(P-value) < 50 and abs(log_2_(fold change)) > 1 informed the selection of peaks relevant for discrimination of tissues. The ability of the peaks to discriminate tissues was then confirmed by visual inspection of their intensity maps across different tissue regions.

### SEM imaging

SEM images of the tissue replicas were acquired with the in-lens detector using a Karl Zeiss XB1540 SEM at 10 kV, after 10 nm Au sputter coating to avoid charging effects.

### Multi-step preprocessing pipeline for machine learning analysis

We developed a multi step preprocessing pipeline to create a common and meaningful representation of a molecular signature across tissue sections and replicas. The pipeline consists of tumour segmentation, spectras correction, normalisation and quantization (Fig S7).

#### Normalisation and quantization

are used to adjust the individual mass spectra in order to remove spectra-to-spectra variability within a given section/replica, generating a common representation with reduced dimensionality that is comparable across all sections/replicas. Each spectra was normalised using the total ion count normalisation method ^32^ normalisation method total ion count (TIC). The normalisation method used in earlier sections using selection of 1000 peaks was not suitable for machine learning analysis because the peaks selection process is based on the full dataset andwhich is inherently leadsing to data contamination. Quantization is frequently used in mass spectra data analysis ^33^. In this study we used equal width binning with bin widths of m/z 0.0125 units. All m/z values and the corresponding intensity values within the bin are represented by the bin centre m/z value and the intensity sum. We assume that the peaks do not slide from one bin to the other during the acquisition, causing errors in peak alignment. This is a reasonable assumption as the mass resolution of the data collected is m/z 0.0250 units which is significantly higher than the selected bin width. However, the m/z values corresponding to ions can be identified more accurately by reducing the bin size. This process leads to a spectra representation of size 92,000 dimensions.

#### Tissue segmentation

is the process of separating the tissue spectras from background (non-tissue) spectras. Due to the high dimensionality of mass spectrometric data we created a single channel representative image using the accumulation of the three representative channels m/z 794.5, m/z 834.5, and m/z 888.6 that are associated with grey matter, white matter and tumour respectively^18^. Using the average threshold we segmented the representative image to use as a mask for the separation of tissue and background spectras. For better segmentation and removal of salt and pepper noise we used median smoothing with a filter size of three.

#### Spectras correction according to the background

(*background correction*) is the process of removing background signals that do not represent the actual tumour signal. Given a spectra in an image we applied Z-Score normalisation using the mean and standard deviation of background spectras of the image. This ensures that the background signal has a mean of zero and standard deviation of one. This correction was applied after observing a high Pearson correlation measurement between tissue and background spectras both in tissue sections (Fig. S8, row 1, middle) and replicas (Fig. S8, row 3, middle) indicating a shared background signal. We can see that this high correlation was fixed after the correction in both tissue sections (Fig. S8, row 2, middle) and replica (Fig. S8, row 4, middle) and did not affect the correlation between tissue and tissue in both tissue sections (Fig. S8, row 2, left) and replica (Fig. S8, row 4, left).

#### Correlation analysis

Correlation-based analysis for every section-replica pair was done by evaluating the Pearson correlation between their mean intensities spectra in the complete mass range m/z 50-1200. This analysis produced a matrix of size nxm where n is the number of sections and m is the number of replicas. Bin (i,j) in this matrix indicates the correlation between section i and replica j. For each replica, we ranked all sections according to their correlation to quantitatively assess whether a replica is most similar to its matched section. To determine whether multiple replicas from the same section maintained their molecular signature, we measured the correlation between each of the replicas of the same tissue with their reference section versus all other sections

### Section-Replica Disease State Classification Architecture

Classification of tissue sections and replicas into high and low grade gliomas was based on a spectra-wise (i.e., pixel-wise) classification in the mass range 600-900 (lipids), followed by majority voting of the spectra classification with a threshold of 0.5. Tissue section and replica classification models were trained and tested independently. We trained multiple classification models, where each model was trained with all sections/replicas excluding sections/replicas that were imaged in the same session or that belong to the same patient. All training spectras were pooled together and then partitioned to train (80%) and validation (20%) sets to assess convergence. Each model was then assessed on the sections/replicas that were not included in its training. Multiple rounds of train-test provided predictions for assessment of the full dataset. The deep neural network architecture was a fully connected neural network consisting of an input layer, three blocks, and an output layer. Each block consisted of the following layers: a dense layer, leaky ReLU activation layer with alpha value of 0.2, batch normalisation layer and a dropout layer with dropout rate of 0.3. Sigmoid activation was used in the output layer to retrieve probability of high/low grade. To avoid over confidence in prediction we applied label smoothing which is a regularisation technique that introduces noise for the labels with a factor value of 0.1. We corrected class imbalance by adjusting the class weights to avoid data imbalance in cross validation. In order to improve convergence, spectras were normalised using min-max technique. This network was trained and implemented using the open source deep learning library of Keras ^34^ and running on Tensorflow ^35^. Training was performed on a high-performance PC workstation equipped with an Intel i7-11700KF 3.60GHz processor, Nvidia GeForce RTX 3070Ti GPU, 64 GB of RAM with a total running time of about 540 minutes for all the models and 30 minutes for a single model.

### Data Visualisation and Statistical Analysis

The bar graphs, volcano plots and heatmaps were prepared using GraphPad Prism 9 (GraphPad Inc., La Jolla, CA, USA). Data in the bar graph is reported as average with standard error of the mean; individual datapoints are also displayed. The schematics in Fig. 1 were prepared using Biorender (biorender.com, 2023). Comparison of total amount of biomolecules between multiple groups was performed by two way analysis of variance (ANOVA). P < 0.05 was considered statistically significant. Comparison between area of replicas with flat vs nanoneedles and comparison between number of features were performed by student t-test. P < 0.05 was considered statistically significant. The p-values used in the volcano plots were calculated by performing multiple unpaired t-test with Holm-Šidák method as implemented in GraphPad Prism 9 (GraphPad Inc., La Jolla, CA, USA).

### Software and Data Availability

The datasets generated and the source code will be made publicly available alongside the publication.

## Author Contributions

**DAM:** Investigation, Formal Analysis, Methodology, Project administration, Software, Validation, Visualization, Writing – original draft; **LAR:** Data Curation, Formal Analysis, Software, Validation, Visualization, Writing – original draft; **NR:** Resources; **CG:** Investigation, Formal Analysis; **VC:** Investigation; **CH:** Resources, **MJ:** Resources, software; **AS:** Resources, Writing – review and editing; **KAJ:** Resources, Writing – review and editing; **MB:** Methdology, Software, Resources, Writing – review and editing; **PB:** Tissue Collection, Data Curation, Formal Analysis, Resources, Writing – review and editing; **AZ:** Formal Analysis, Funding acquisition, Investigation, Methodology, Software, Validation, Visualization, Writing – original draft; **CC:** Conceptualization, Funding acquisition, Methodology, Project administration, Supervision, Visualization, Writing – original draft.

## Acknowledgements

CC acknowledges funding from the European Union under the ERC Starting Grant ENBION 759577; the Wellcome Leap Delta Tissue Programme; and Debra UK. PB acknowledges the contribution of the Lothian NRS bioresource. CG acknowledges King’s-China Scholarship Council PhD Scholarship and the London Centre for Nanotechnology. AZ acknowledges funding from the Wellcome Leap Delta Tissue Programme.

