## Supplementary Information for "Nondestructive Spatial Lipidomics for Glioma Classification"

### Materials and Methods

#### COMSOL finite element simulations

The finite element simulations were performed by using COMSOL Multiphysics® 5.4 and Heat Transfer module. Two geometries were defined: a rectangle 525 µm wide and 8 mm tall to simulate the nN chip and a rectangle 19.475 mm wide, 4 mm / 8 mm tall, to simulate a tissue specimen with smaller or larger contact area with the nN chip, respectively. A triangular 0.6 mm mesh was defined for both nN chip and tissue. The material properties were selected from the COMSOL library: silicon for the nN chip and water/ice for the tissue. The initial temperature of the nN chip was 25 °C, the initial temperature of the tissue was -20 °C. The side of the tissue further from the nN chip was kept constant at -20 °C, at all times during the simulation. An insulating layer was defined around all the other sides of the objects but the one where they were in contact. A time range of 1 s and step of 0.05 s were used for the simulation.

### Supplementary Information

#### Principal Component Analysis

We used principal component analysis (PCA) to perform an internal validation of the clustering analysis, by assessing the DESI-MS spectral variance between and within clusters. Mapping the distribution of principal component scores also provided further insight into the heterogeneity within the tumour cluster. The first principal component, PC 1 accounted for the 68 % of the variance, arising from the variation of the main spectral profile. The second and third principal components, PC 2 (6.8 %) and PC 3 (4.6 %) accounted for a large proportion of the variance within the the dataset (Fig. S3-a). The loading from PC 2 showed positive peaks associated to white matter, in particular the sulfatide peak at 888.6 m/z and phosphatidylserine at 788.5 m/z [1]. The negative peaks were mainly associated to grey matter and tumour, in particular the phosphatidylserine peak at 834.5 m/z (grey matter), the phosphatidylethanolamine peak at 766.5 m/z (tumour) and the phosphatidylinositol peak at 885.5 m/z (grey matter and tumour) [1,2]. The loadings from PC 3 showed positive peaks associated to grey matter at 600.5 m/z (putatively Cer), 654.5 m/z (putative Cer), 746.5 m/z (putatively PC/PE), 774.5 m/z (putatively PC/PE), 790.5 m/z (putatively PC/PE), 834.5 m/z (putatively PS) and negative peaks associated to the tumour at 611.5 m/z (putatively phosphatidylglycerol, PG), 716 m/z (putatively PS/PE), 744.5 m/z (putatively PS/PE), 764.5 m/z (putatively PE), 788.5 m/z (putatively PS), 812.5 m/z (putatively PC), 885.5 m/z (putatively PI). The score maps for PC 2 and PC 3 highlighted the association of the loading of individual components to the different tissues (Fig. S3-b). The map of PC 2 scores distinguished white matter from grey matter and tumour. The map of PC 3 scores distinguished the tumour against grey matter and white matter.

Principal component analysis in combination with the assigned pixel identity from hierarchical cluster analysis provided further insight into the inherent spectral variability of tissue section and replica (Fig. S3-c). For white and grey matter, the data from the replica showed a broadening of distribution and a shift towards null score. The tumour cluster instead remained more similarly distributed between section and replica. On both section and replica, PC 3 distinguished between tumour and grey matter, with the white matter in between. The grey matter distribution of PC 3-scores showed a bimodal distribution. The white matter cluster in the replica showed a shift towards the grey matter. In both section and replica, the tumour cluster showed a sharp distribution with a tail towards the other clusters. Spatially, these tails localised to the margin region between tumour and healthy tissue both in the section (Fig. S3-d) and in the replica (Fig. S3-e), suggesting an ability of PCA analysis to map the margin region of the tumour and the transitions between tissues.

The principal component analysis indicates that the information contained within the data from the replica broadly aligns with that of the original sample. Moreover, the intracluster variance in PCA loadings can map tumour margins and transitions between tissues in the replica as well as in the section.

#### Supplementary Figures and Tables

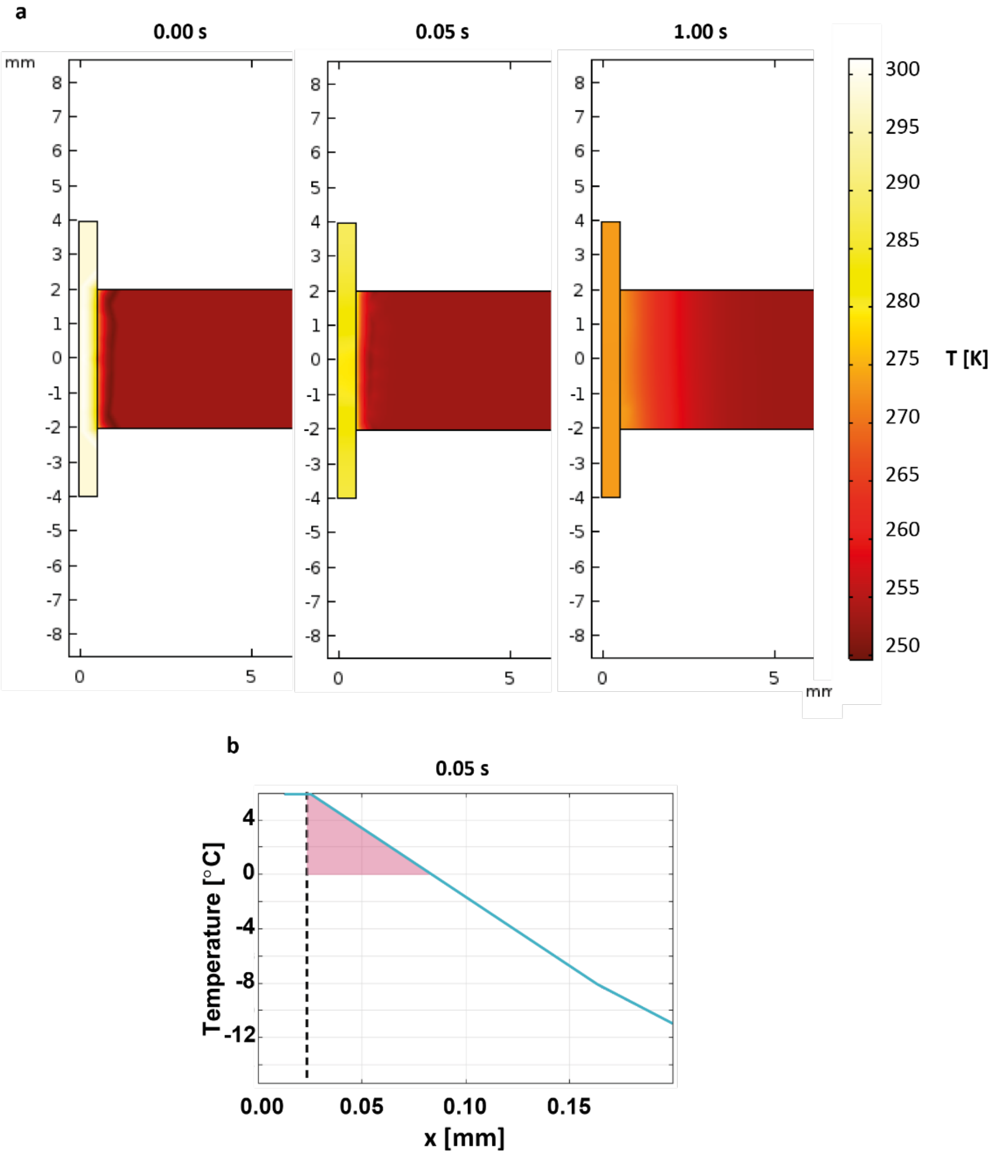

**Figure S1: Computational modelling of tissue interfacing.** (a) Cross-sectional snapshots of the simulation of heat transfer between chip and tissue. The chip is the tall and narrow element on the left while the tissue is the longer element on the right. From left to right the images show the distribution of the temperature at 0 s, 0.05 s and 1 s of the simulation. The chip was initially at 25 °C = 298.15 K, while the specimen was at -20 °C = 253.15 K. In the first 0.05 s, at the interface with the chip, the temperature of the specimen rose above 0 °C = 273.15 K. At 1 s the specimen re-froze onto the chip. (b) Plot of the temperature variation along a line drawn from the chip to the specimen, at 0.05 s. The dashed line marks the interface between chip and specimen. The area highlighted in pink shows the thawed region of ~60 µm within the tissue after 0.05 s.

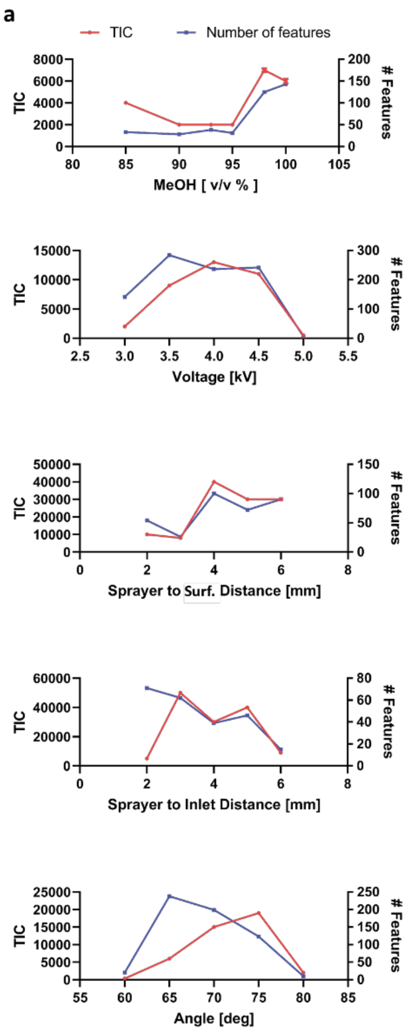

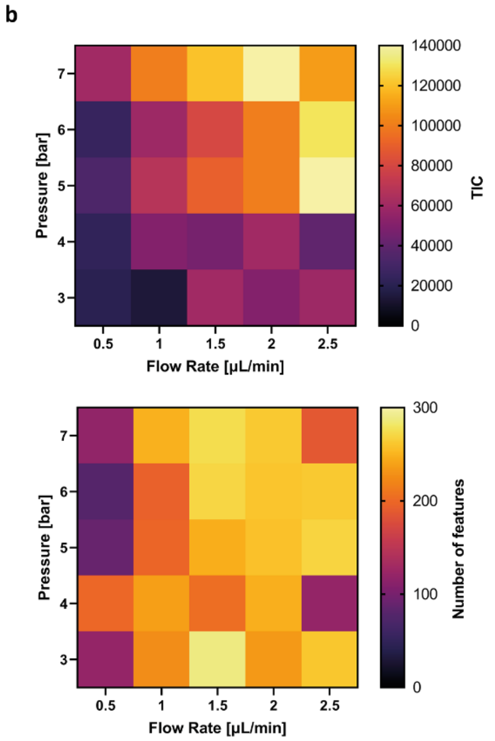

c

| Parameters | Std | Opt |
|---|---|---|
| [MeOH] | 100 % | 100 % |
| Voltage | 4.5 kV | 4 kV |
| S-I distance | 2 mm | 2 mm |
| S-S distance | 2 mm | 4 mm |
| S-S Angle | 60° | 65° |
| Pressure - flow rate | 5 bar - 2 µL/min | 5 bar - 2.5 µL/min |

**Figure S2: Optimisation of DESI settings for imaging.** Parameters optimised to maximise Total Ion Count (TIC) and number of features after denoising. (a) Line plots, showing TIC and number of features as a function of the optimised parameters: MeOH : $H_2O$ solvent composition, voltage, Sprayer to Surface distance, Sprayer to Inlet distance, Sprayer to Surface angle. (b) Heatmaps showing TIC (top) and number of features as a function of the inter-related gas pressure and solvent flow rate. (c) Table showing the comparison between the standard and optimised settings.

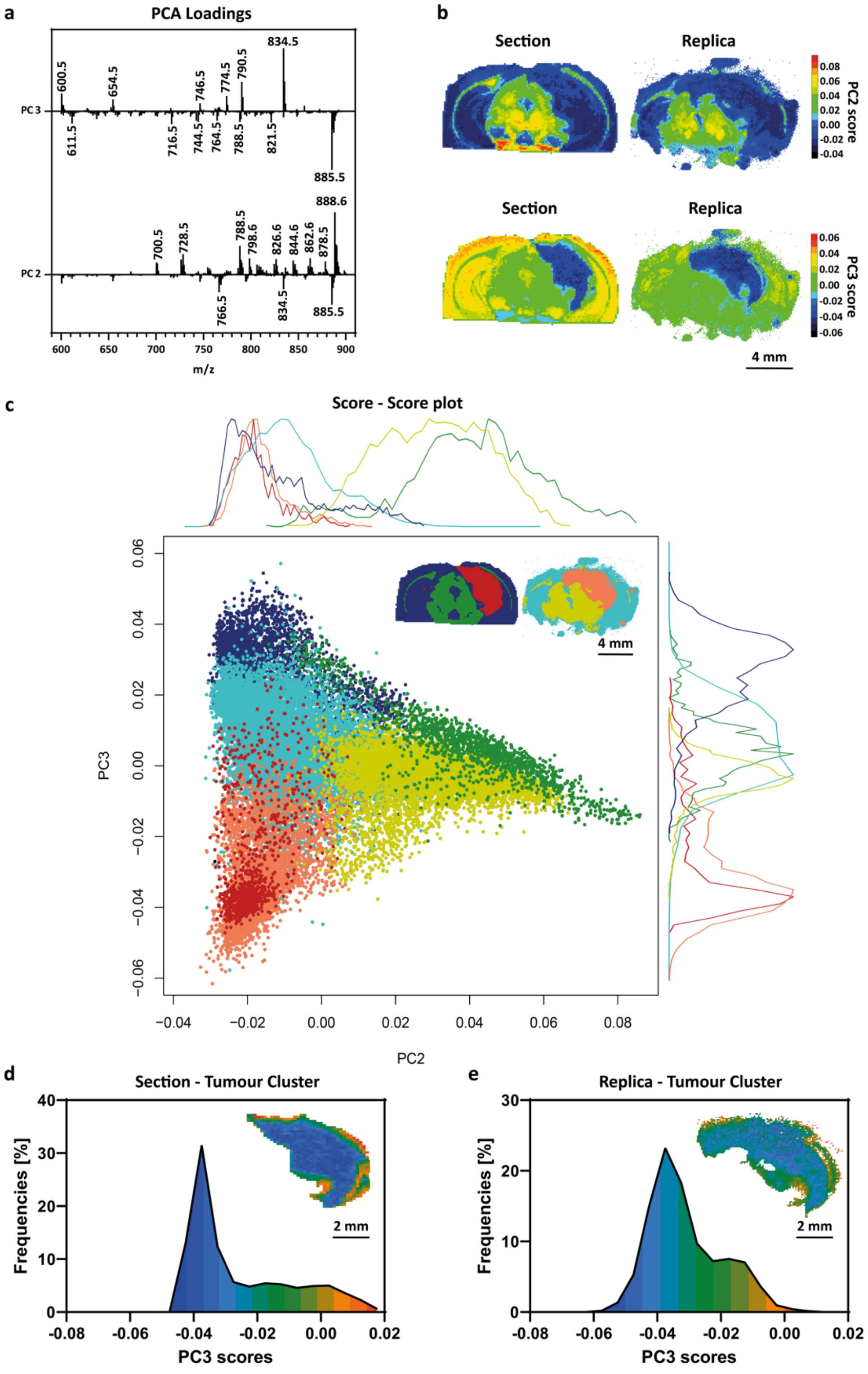

**Figure S3: Principal Component Analysis.** (a) Loadings of the principal components PC2 and PC3, obtained by principal component analysis (PCA) of DESI-MS images acquired from matching section and replica of tumour-bearing murine brain. (b) Maps of the scores from PCA for PC2 and PC3, showing that PC2 discriminates white matter vs the other 2 tissues, while PC3 discriminates healthy tissue vs tumour. (c) Score-score plot, showing the score value for every spectrum in section and replica. The colours were associated to the clusters as obtained by HCA and spatially distributed as shown in the inset cluster maps, to allow the observation of inter- and intra-cluster spectral variance. Along the axis, graphs representing spectra frequency distributions vs score of the principal component, for each cluster associated to grey matter/white matter/tumour and section/replica. (d) Graph representing the spectral frequency distribution of PC3 scores in the tumour-associated cluster in the section, with colours showing the spectra spatial location and associating respectively the lower and higher score values to core and peripheral spatial location in the inset map. (e) Graph representing the spectral frequency distribution of PC3 scores in the tumour-associated cluster in the replica, with colours associating respectively the lower and higher score values to core and peripheral spatial location in the inset map, in the replica as well as observed in the section.

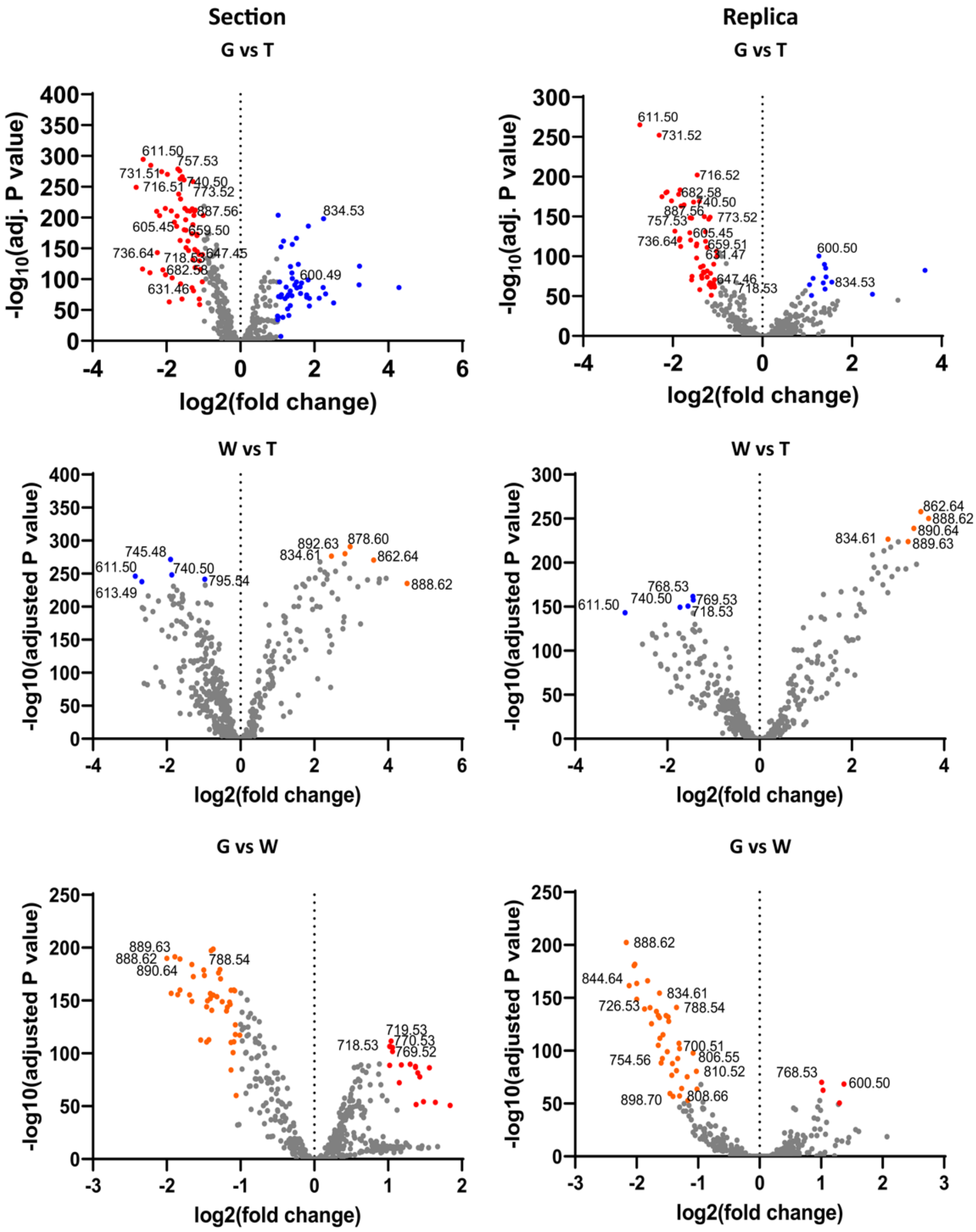

**Figure S4: Differential lipid abundance across grey, white matter and tumour.** Volcano plots showing the differential lipid distribution in pairwise comparisons between grey matter, white matter and tumour for section and replica, with highlighted representative lipid peaks. The volcano plots informed the selection of the 33 most representative peaks used in Fig. 4 j, k. which are labelled on the plots.

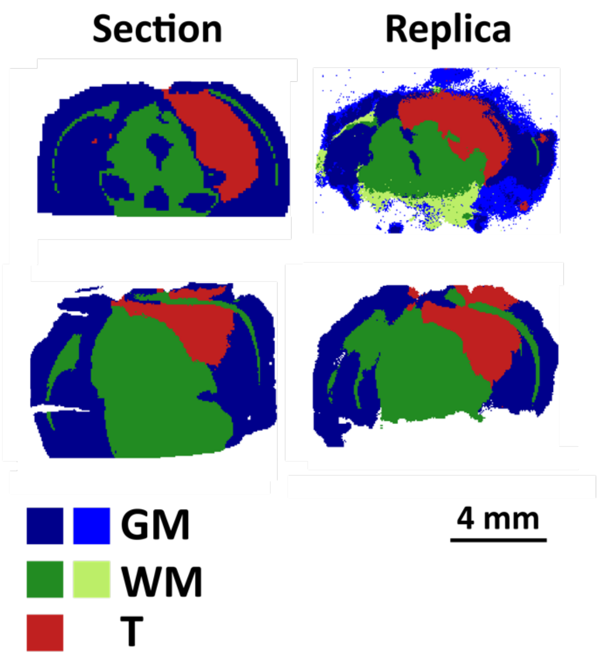

**Figure S5: Hierarchical cluster analysis of tumour-bearing mouse brains.** Maps of clusters associated to grey, white matter and tumour, obtained by HCA of DESI-MSI datasets of brain section and replica for two different tumour-bearing mice, supporting the reproducibility of the unsupervised classification of the tissue in section and replica.

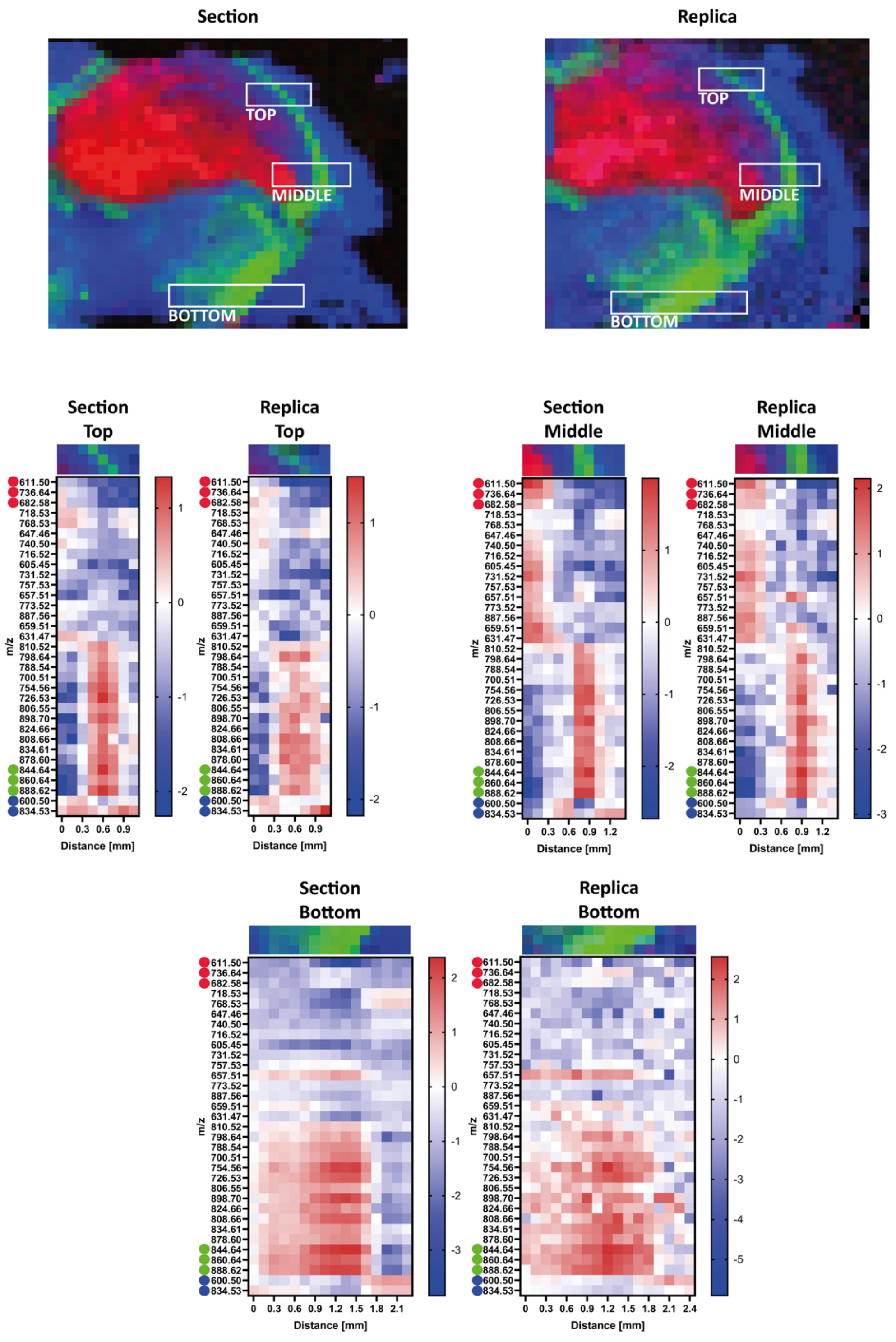

**Figure S6: Comparison of spatial pattern and lipid abundance across multiple regions.** Heatmaps showing the relative abundance along the major axis of the white boxes of the 33 lipid peaks identified by differential analysis within the section and the replica. Matching pattern and lipid abundance are shown in region of the corpus callosum near the tumour (TOP and MIDDLE) and in the region of the peduncle (BOTTOM).

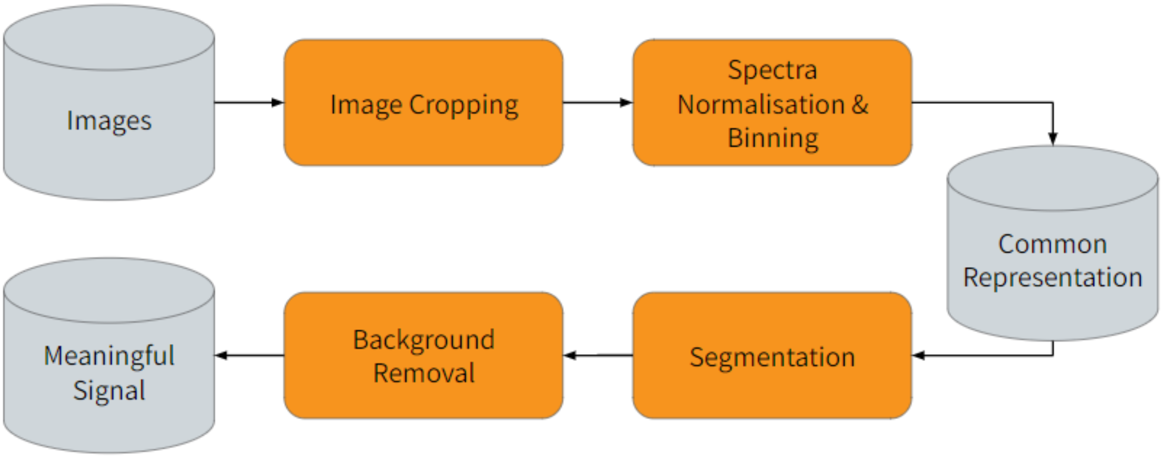

**Figure S7: Multistep preprocessing pipeline.** Schematic representation of the multistep preprocessing pipeline used to support the deep neural network analysis.

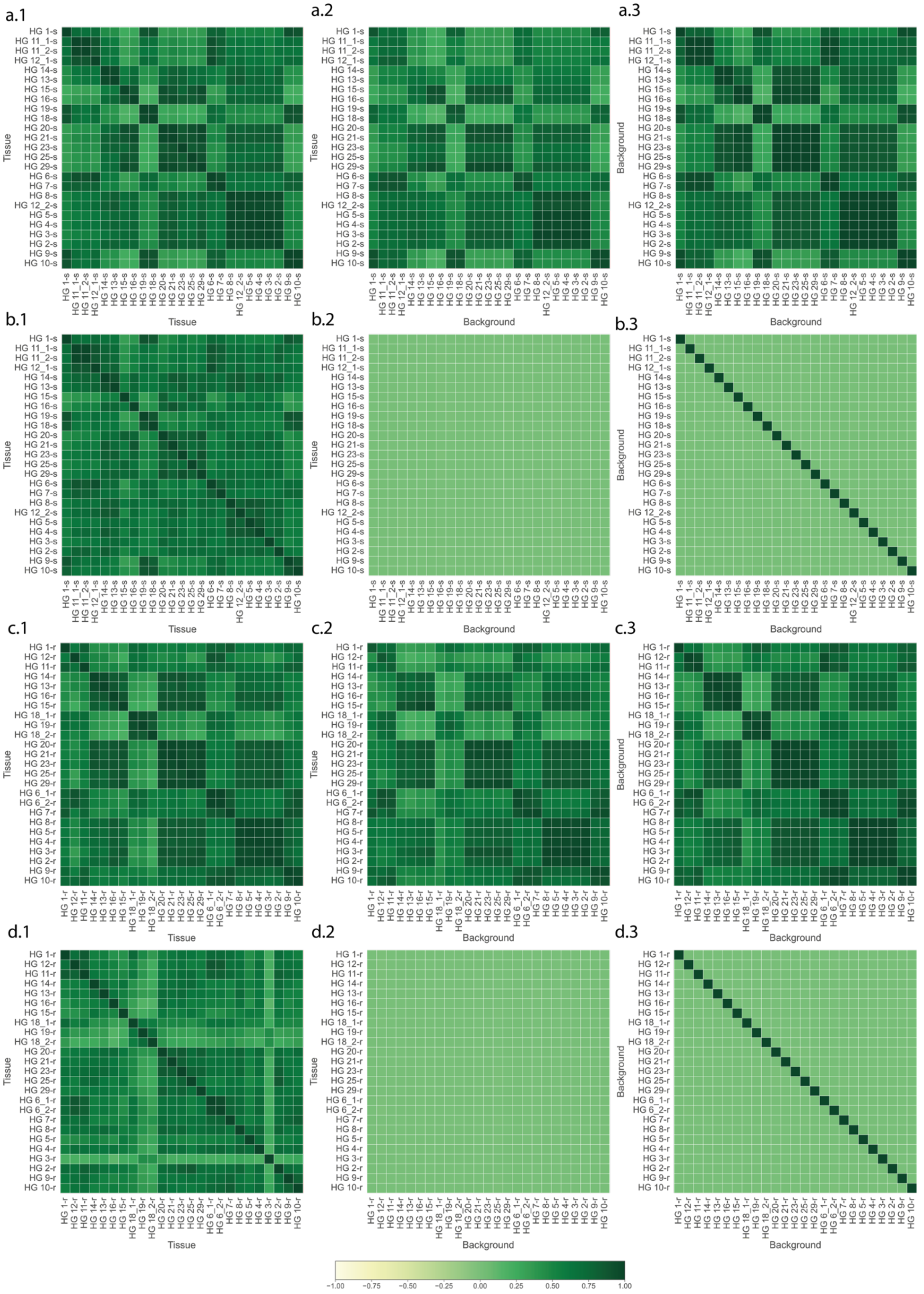

**Figure S8: Similarity between all tissue and background permutations in both tissue sections and nanobiopsy replicas using Pearson correlation.** (First row) Heatmap of Pearson's correlation values between tissue and tissue, tissue and background, background and background in tissue sections before correction. (Second row) Heatmap of correlation values between tissue and tissue, tissue and background, background and background in tissue sections after correction. (Third row) Heatmap of correlation values between tissue and tissue, tissue and background, background and background in nanobiopsy replicas before correction. (Fourth row) Heatmap of correlation values between tissue and tissue, tissue and background, background and background in molecular replicas after correction.

| Biopsy | Sample Label | Sample Type | histopathological annotations | WHO Grade | DNN Classification |
|---|---|---|---|---|---|
| 1 | HG 1-r | replica | oligodendroglioma, 1p19 q codel | II | LOW |
| 1 | HG 1-s | section | oligodendroglioma, 1p19 q codel | II | LOW |
| 2 | HG 2-s | section | oligodendroglioma | II | HIGH |
| 2 | HG 2-r | replica | oligodendroglioma | II | LOW |
| 3 | HG 3-s | section | glioblastoma multiforme | IV | HIGH |
| 3 | HG 3-r | replica | glioblastoma multiforme | IV | LOW |
| 4 | HG 4-s | section | glioblastoma multiforme | IV | HIGH |
| 4 | HG 4-r | replica | glioblastoma multiforme | IV | HIGH |
| 5 | HG 5-s | section | glioblastoma multiforme | IV | HIGH |
| 5 | HG 5-r | replica | glioblastoma multiforme | IV | HIGH |
| 6 | HG 6_1-r | replica | glioblastoma multiforme | IV | HIGH |
| 6 | HG 6_2-r | replica | glioblastoma multiforme | IV | HIGH |
| 6 | HG 6-s | section | glioblastoma multiforme | IV | HIGH |
| 7 | HG 7-r | replica | glioblastoma multiforme | IV | HIGH |
| 7 | HG 7-s | section | glioblastoma multiforme | IV | HIGH |
| 8 | HG 8-s | section | anaplastic astrocytoma | III | HIGH |
| 8 | HG 8-r | replica | anaplastic astrocytoma | III | LOW |
| 9 | HG 9-r | replica | anaplastic oligodendroglioma, 1p19q codel | III | HIGH |
| 9 | HG 9-s | section | anaplastic oligodendroglioma, 1p19q codel | III | LOW |
| 10 | HG 10-r | replica | secondary GBM – transf. of oligoastrocytoma | IV | HIGH |
| 10 | HG 10-s | section | secondary GBM – transf. of oligoastrocytoma | IV | LOW |
| 11 | HG 11_1-s | section | anaplastic astrocytoma | III | HIGH |
| 11 | HG 11_2-s | section | anaplastic astrocytoma | III | HIGH |
| 11 | HG 11-r | replica | anaplastic astrocytoma | III | HIGH |
| 12 | HG 12_1-s | section | glioblastoma multiforme | IV | HIGH |
| 12 | HG 12-r | replica | glioblastoma multiforme | IV | HIGH |
| 12 | HG 12_2-s | section | glioblastoma multiforme | IV | HIGH |
| 13 | HG 13-r | replica | glioblastoma multiforme | IV | HIGH |
| 13 | HG 13-s | section | glioblastoma multiforme | IV | HIGH |
| 14 | HG 14-r | replica | glioblastoma multiforme | IV | HIGH |
| 14 | HG 14-s | section | glioblastoma multiforme | IV | HIGH |
| 15 | HG 15-r | replica | glioblastoma multiforme | IV | HIGH |
| 15 | HG 15-s | section | glioblastoma multiforme | IV | LOW |
| 16 | HG 16-r | replica | pilocytic astroctytoma | I | HIGH |
| 16 | HG 16-s | section | pilocytic astroctytoma | I | HIGH |
| 18 | HG 18_1-r | replica | diffuse astrocytoma | II | HIGH |
| 18 | HG 18_2-r | replica | diffuse astrocytoma | II | HIGH |
| 18 | HG 18-s | section | diffuse astrocytoma | II | HIGH |
| 19 | HG 19-r | replica | oligodendorglioma | II | HIGH |
| 19 | HG 19-s | section | oligodendorglioma | II | LOW |
| 20 | HG 20-r | replica | diffuse astrocytoma, IDH1 mutant | II | LOW |

| | | | | | |
|---|---|---|---|---|---|
| 20 | HG 20-s | section | diffuse astrocytoma, IDH1 mutant | II | HIGH |
| 21 | HG 21-r | replica | anaplastic oligodendroglioma, 1p19q codel, IDH1 mutant | III | LOW |
| 21 | HG 21-s | section | anaplastic oligodendroglioma, 1p19q codel, IDH1 mutant | III | LOW |
| 23 | HG 23-r | replica | pilocytic astrocytoma | I | LOW |
| 23 | HG 23-s | section | pilocytic astrocytoma | I | LOW |
| 25 | HG 25-r | replica | oligodendroglioma, 1p19 q codel, IDH1 mutant | II | LOW |
| 25 | HG 25-s | section | oligodendroglioma, 1p19 q codel, IDH1 mutant | II | LOW |
| 29 | HG 29-r | replica | oligodendroglioma, 1p19q codel | II | LOW |
| 29 | HG 29-s | section | oligodendroglioma, 1p19q codel | II | LOW |

**Table S1: Summary table of analysed samples.** The table indicates the unique biopsy number, the label used to identify each sample in the manuscript text and figures, whether the sample is a tissue section or a molecular replica, the histopathological annotations of the biopsy, the WHO grade and the results of the deep neural network classification colour coded in green for correct assignments and orange for mis-assignments.

| Train samples | Test samples | Num. train spectras | Num. validation spectras | Num test spectras |
|---|---|---|---|---|
| HG 11_1-s, HG 11_2-s, HG 12_1-s, HG 14-s, HG 13-s, HG 15-s, HG 16-s, HG 19-s, HG 18-s, HG 20-s, HG 21-s, HG 23-s, HG 25-s, HG 29-s, HG 6-s, HG 7-s, HG 8-s, HG 12_2-s, HG 5-s, HG 4-s, HG 3-s, HG 2-s, HG 9-s, HG 10-s | HG 1-s | 24206 | 6052 | 1704 |
| HG 14-s, HG 13-s, HG 15-s, HG 16-s, HG 19-s, HG 18-s, HG 20-s, HG 21-s, HG 23-s, HG 25-s, HG 29-s, HG 6-s, HG 7-s, HG 8-s, HG 5-s, HG 4-s, HG 3-s, HG 2-s, HG 9-s, HG 10-s, HG 1-s | HG 11_1-s, HG 11_2-s, HG 12_1-s, HG 12_2-s | 20770 | 5193 | 5999 |
| HG 11_1-s, HG 11_2-s, HG 12_1-s, HG 15-s, HG 16-s, HG 19-s, HG 18-s, HG 20-s, HG 21-s, HG 23-s, HG 25-s, HG 29-s, HG 6-s, HG 7-s, HG 8-s, HG 12_2-s, HG 5-s, HG 4-s, HG 3-s, HG 2-s, HG 9-s, HG 10-s, HG 1-s | HG 14-s, HG 13-s | 23054 | 5764 | 3144 |
| HG 11_1-s, HG 11_2-s, HG 12_1-s, HG 14-s, HG 13-s, HG 19-s, HG 18-s, HG 20-s, HG 21-s, HG 23-s, HG 25-s, HG 29-s, HG 6-s, HG 7-s, HG 8-s, HG 12_2-s, HG 5-s, HG 4-s, HG 3-s, HG 2-s, HG 9-s, HG 10-s, HG 1-s | HG 15-s, HG 16-s | 23575 | 5894 | 2493 |
| HG 11_1-s, HG 11_2-s, HG 12_1-s, HG 14-s, HG 13-s, HG 15-s, HG 16-s, HG 20-s, HG 21-s, HG 23-s, HG 25-s, HG 29-s, HG 6-s, HG 7-s, HG 8-s, HG 12_2-s, HG 5-s, HG 4-s, HG 3-s, HG 2-s, HG 9-s, HG 10-s, HG 1-s | HG 19-s, HG 18-s | 24029 | 6008 | 1925 |
| HG 11_1-s, HG 11_2-s, HG 12_1-s, HG 14-s, HG 13-s, HG 15-s, HG 16-s, HG 19-s, HG 18-s, HG 6-s, HG 7-s, HG 8-s, HG 12_2-s, HG 5-s, HG 4-s, HG 3-s, HG 2-s, HG 9-s, HG 10-s, HG 1-s | HG 20-s, HG 21-s, HG 23-s, HG 25-s, HG 29-s | 19195 | 4799 | 7968 |
| HG 11_1-s, HG 11_2-s, HG 12_1-s, HG 14-s, HG 13-s, HG 15-s, HG 16-s, HG 19-s, HG 18-s, HG 20-s, HG 21-s, HG 23-s, HG 25-s, HG 29-s, HG 8-s, HG 12_2-s, HG 5-s, HG 4-s, HG 3-s, HG 2-s, HG 9-s, HG 10-s, HG 1-s | HG 6-s, HG 7-s | 24432 | 6109 | 1421 |
| HG 11_1-s, HG 11_2-s, HG 14-s, HG 13-s, HG 15-s, HG 16-s, HG 19-s, HG 18-s, HG 20-s, HG 21-s, HG 23-s, HG 25-s, HG 29-s, HG 6-s, HG 7-s, HG 9-s, HG 10-s, HG 1-s | HG 12_1-s, HG 8-s, HG 12_2-s, HG 5-s, HG 4-s, HG 3-s, HG 2-s | 18332 | 4584 | 9046 |

| | | | | |
|---|---|---|---|---|
| HG 11_1-s, HG 11_2-s, HG 12_1-s, HG 14-s, HG 13-s, HG 15-s, HG 16-s, HG 19-s, HG 18-s, HG 20-s, HG 21-s, HG 23-s, HG 25-s, HG 29-s, HG 6-s, HG 7-s, HG 8-s, HG 12_2-s, HG 5-s, HG 4-s, HG 3-s, HG 2-s, HG 1-s | HG 9-s, HG 10-s | 24248 | 6063 | 1651 |
| HG 12-r, HG 11-r, HG 14-r, HG 13-r, HG 16-r, HG 15-r, HG 18_1-r, HG 19-r, HG 18_2-r, HG 20-r, HG 21-r, HG 23-r, HG 25-r, HG 29-r, HG 6_1-r, HG 6_2-r, HG 7-r, HG 8-r, HG 5-r, HG 4-r, HG 3-r, HG 2-r, HG 9-r, HG 10-r | HG 1-r | 34952 | 8738 | 2543 |
| HG 14-r, HG 13-r, HG 16-r, HG 15-r, HG 18_1-r, HG 19-r, HG 18_2-r, HG 20-r, HG 21-r, HG 23-r, HG 25-r, HG 29-r, HG 6_1-r, HG 6_2-r, HG 7-r, HG 8-r, HG 5-r, HG 4-r, HG 3-r, HG 2-r, HG 9-r, HG 10-r, HG 1-r | HG 12-r, HG 11-r | 35132 | 8784 | 2317 |
| HG 12-r, HG 11-r, HG 16-r, HG 15-r, HG 18_1-r, HG 19-r, HG 18_2-r, HG 20-r, HG 21-r, HG 23-r, HG 25-r, HG 29-r, HG 6_1-r, HG 6_2-r, HG 7-r, HG 8-r, HG 5-r, HG 4-r, HG 3-r, HG 2-r, HG 9-r, HG 10-r, HG 1-r | HG 14-r, HG 13-r | 32926 | 8232 | 5075 |
| HG 12-r, HG 11-r, HG 14-r, HG 13-r, HG 18_1-r, HG 19-r, HG 18_2-r, HG 20-r, HG 21-r, HG 23-r, HG 25-r, HG 29-r, HG 6_1-r, HG 6_2-r, HG 7-r, HG 8-r, HG 5-r, HG 4-r, HG 3-r, HG 2-r, HG 9-r, HG 10-r, HG 1-r | HG 16-r, HG 15-r | 34492 | 8623 | 3118 |
| HG 12-r, HG 11-r, HG 14-r, HG 13-r, HG 16-r, HG 15-r, HG 20-r, HG 21-r, HG 23-r, HG 25-r, HG 29-r, HG 6_1-r, HG 6_2-r, HG 7-r, HG 8-r, HG 5-r, HG 4-r, HG 3-r, HG 2-r, HG 9-r, HG 10-r, HG 1-r | HG 18_1-r, HG 19-r, HG 18_2-r | 31299 | 7825 | 7109 |
| HG 12-r, HG 11-r, HG 14-r, HG 13-r, HG 16-r, HG 15-r, HG 18_1-r, HG 19-r, HG 18_2-r, HG 6_1-r, HG 6_2-r, HG 7-r, HG 8-r, HG 5-r, HG 4-r, HG 3-r, HG 2-r, HG 9-r, HG 10-r, HG 1-r | HG 20-r, HG 21-r, HG 23-r, HG 25-r, HG 29-r | 29704 | 7426 | 9103 |
| HG 12-r, HG 11-r, HG 14-r, HG 13-r, HG 16-r, HG 15-r, HG 18_1-r, HG 19-r, HG 18_2-r, HG 20-r, HG 21-r, HG 23-r, HG 25-r, HG 29-r, HG 8-r, HG 5-r, HG 4-r, HG 3-r, HG 2-r, HG 9-r, HG 10-r, HG 1-r | HG 6_1-r, HG 6_2-r, HG 7-r | 34564 | 8642 | 3027 |

| | | | | |
|---|---|---|---|---|
| HG 12-r, HG 11-r, HG 14-r, HG 13-r, HG 16-r, HG 15-r, HG 18_1-r, HG 19-r, HG 18_2-r, HG 20-r, HG 21-r, HG 23-r, HG 25-r, HG 29-r, HG 6_1-r, HG 6_2-r, HG 7-r, HG 9-r, HG 10-r, HG 1-r | HG 8-r, HG 5-r, HG 4-r, HG 3-r, HG 2-r | 28989 | 7248 | 9996 |
| HG 12-r, HG 11-r, HG 14-r, HG 13-r, HG 16-r, HG 15-r, HG 18_1-r, HG 19-r, HG 18_2-r, HG 20-r, HG 21-r, HG 23-r, HG 25-r, HG 29-r, HG 6_1-r, HG 6_2-r, HG 7-r, HG 8-r, HG 5-r, HG 4-r, HG 3-r, HG 2-r, HG 1-r | HG 9-r, HG 10-r | 33830 | 8458 | 3945 |

**Table S1: leave one batch and patient out training process.** Table showing number of train, validation and test spectra as a function of the combination of train and test dataset.

| Train data sample type | Test data sample type | Balanced accuracy | AUC | Misclassified samples |
|---|---|---|---|---|
| section tissue | section tissue | 0.71 | 0.8 | HG 10-s, HG 15-s, HG 16-s, HG 2-s, HG 20-s, HG 21-s, HG 9-s |
| section tissue | replica tissue | 0.6 | 0.7 | HG 1-r, HG 16-r, HG 18_1-r, HG 18_2-r, HG 19-r, HG 2-r, HG 20-r, HG 29-r |
| replica tissue | replica tissue | 0.7 | 0.8 | HG 16-r, HG 18_1-r, HG 18_2-r, HG 19-r, HG 21-r, HG 3-r, HG 8-r |
| replica tissue | section tissue | 0.51 | 0.6 | HG 10-s, HG 13-s, HG 14-s, HG 16-s, HG 2-s, HG 20-s, HG 23-s, HG 25-s, HG 29-s, HG 8-s, HG 9-s |

**Table S3: Performance metrics for glioma classification.** Table showing balanced accuracies, AUC and misclassified samples as a function of the combination of train and test dataset.
